# New llama-derived nanobodies enable selective detection and capture of Z-DNA

**DOI:** 10.64898/2026.11.17.765674

**Authors:** Mathias I. Nielsen, Peter C. Fridy, Jonah Da Silva, Mohammad Pourshamsi, Laurens Wachsmuth, Jonathan Kittel, Hua Jiang, Bryant D. Miller, Martin S. Taylor, Brian T. Chait, Manolis Pasparakis, J. Michelle Kahlenberg, Tomas Mustelin, Michael P. Rout, John LaCava

## Abstract

Z-DNA is a left-handed DNA conformation that forms transiently at susceptible genomic sequences during processes such as negative supercoiling and active transcription. In the cytoplasm, the innate immune receptor ZBP1 senses Z-form nucleic acids (Z-NAs, which could be DNA or RNA), triggering type I interferon production and programmed cell death. Viral infections, endogenous repetitive-element-derived RNAs, and oxidized mitochondrial DNA can generate Z-NAs, nominating them as ZBP1 agonists. Conversely, ADAR1 is proposed to recognize and edit endogenous Z-RNAs, thereby muting inappropriate innate immune responses. Both proteins engage Z-NAs through Zα domains, which are evolutionarily conserved across vertebrates. Progress in defining Z-NA biology has been limited by near-exclusive reliance on the Z22 monoclonal antibody, the field’s principal detection reagent. To expand the availability of distinct, validated anti-Z-NA affinity reagents, we generated and characterized llama-derived nanobodies. Llamas were immunized with brominated poly(dG:dC) DNA, and four recombinant nanobodies (nZ6, nZ6.2, nZ10, and nZ11) that bound the antigen were identified through affinity enrichment and mass spectrometry-guided discovery. The nanobodies selectively recognized Z-DNA in biochemical assays and displayed nanomolar affinities, while homodimerized nZ6.2 achieved picomolar binding. Unlike Zα domains, nZ6.2 does not induce B-to-Z transitions, indicating recognition of pre-existing Z-DNA and immobilized nZ6.2 selectively co-immunoprecipitated ADAR1 and ZBP1 with Z-DNA but not B-DNA. Furthermore, nZ6.2 detected Z-DNA-related structures in a human keratinocyte cell line and in neutrophils from patients with systemic lupus erythematous, producing staining patterns overlapping those of Z22. These renewable, engineerable binders provide orthogonal reagents for investigating Z-NA biology, interferon signaling, and autoimmune disease.

## Introduction

Z-DNA is a non-canonical, dynamic DNA conformation that can arise under physiological conditions and may play regulatory roles in the genome (1, 2). It presents as an alternative structural form of DNA exhibiting a left-handed, double-helical twist and characteristic zigzag pattern (3). Z-DNA arises most frequently in alternating purine–pyrimidine sequences, especially GC repeats, and it is stabilized by high salt, chemical modifications, negative supercoiling, or active transcription (2, 4–8). *In vitro*, bromination of DNA, typically induced by high salt treatment, is often used to sterically ‘lock’ DNA into the Z-conformation (9). In cells, Z-DNA is thought to form at actively transcribed regions of DNA, it has been implicated in the regulation of gene expression, and was presumed to promote interactions with specific Z-DNA binding proteins (10). Notably, ZBP1 and the p150 isoform of ADAR1, respectively, are the only annotated mammalian Z-DNA binding proteins (reviewed in (11, 12). Although, both ZBP1 and the p150 isoform of ADAR1 were initially characterized as Z-DNA binding proteins *in vitro* (*13, 14*), the majority of recent evidence points towards their major biological functions occurring in the cytoplasm, where they bind Z-DNA or Z-RNA (Z-NA)(11, 12, 15), of viral or endogenous origins, and trigger innate immune signalling cascades (11, 16–18). Indeed, the E3L protein of vaccinia virus confers interferon resistance to the virus by mitigating ZBP1-derived immune signaling; E3L binds and sequesters vaccinia Z-RNAs, mitigating recognition by ZBP1 and consequent anti-viral necroptosis (19). The domains within ZBP1, ADAR1 p150, and vaccinia E3L that mediate Z-NA interactions are known as Zα domains (13–15, 19, 20). Zα domains bind to pre-formed Z-NAs but have also been shown to bind B-form DNA and A-form RNA, albeit more weakly, and induce Z-NA formation by lowering the energetic barrier of B/A to Z transitions (B/A→Z) (21–26).

To facilitate the study of Z-DNA, several monoclonal antibodies were raised (27–31). Early monoclonal antibodies known as Z22, Z44, and D11 were the subjects of numerous biochemical and structural Z-DNA characterizations, proving that Z-DNA exhibits multiple distinctive antigenic structural features. Like the Z⍺ domain, Z-DNA-binding monoclonal antibodies have also exhibited binding to Z-RNA (32) – therefore (unless shown otherwise), these reagents can be considered non-discriminating Z-NA binding reagents. Z-NA-binding antibodies have been shown to promote B/A→Z transitions, albeit less efficiently than Z⍺ domains (31, 33–35). Among the early ⍺-Z-NA monoclonal antibodies, Z22 became the most widely used reagent for detecting Z-NAs in molecular cell biology. This clone is readily available from commercial vendors, while other clones can be more difficult to obtain. Hence, there is effectively only a single, well-characterized, commercially available ⍺-Z-NA reagent in global use for both Z-NA detection and capture.

Our group became interested in Z-NA through our investigations of the underlying causes of Systemic Lupus Erythematosus (SLE) – a disease characterized by chronically elevated type I interferon signaling (36, 37). SLE patients and autoimmune, lupus-prone mice have circulating antibodies that recognize left-handed Z-DNA (29, 30, 38–41); a recent model posits that, in SLE skin photosensitivity, UVB-induced mitochondrial oxidative stress generates cytosolic Z-DNA (derived from oxidized mtDNA), resulting in ZBP1-mediated type I interferon signaling (17). When we began our research into this area, we were uncomfortable with relying upon a single validated reagent for both capture and detection, since this can amplify false positives when there is cross-reactivity (42, 43). Rather than attempting to raise additional monoclonal antibodies based on canonical mammalian IgGs, we raised camelid heavy chain-only-derived nanobodies, a.k.a. V_H_H antibodies (44). In additional to being smaller (∼15 kDa nanobody vs. ∼150 kDa IgG), nanobodies exhibit numerous advantages over traditional antibodies: they are encoded by a single polypeptide, are cheaply and easily produced in bacteria, they exhibit simpler folding, greater stability, and greater solubility, and they are easier to engineer as fusion proteins, making them applicable in a wider array of assays, including intracellular targeting and live cell imaging (44, 45). In addition, when our group became interested in Z-NA, we initially worked with a commercially supplied Z22 monoclonal antibody; however, we experienced significant lot-to-lot variability in performance, which confounded our results and delayed our experimental work.

## Materials and Methods

### Immunization and candidate selection

Following a published nanobody production pipeline (46, 47) and using the same poly-GC antigen as for the development of the Z22 monoclonal antibody (27, 39): animals were immunized with an antigen mixture containing 100 μg of a brominated poly-GC sequence (∼1.1 kb, Amersham Biosciences, product #27-7910-03 [no longer available]) complexed with 100 μg methylated BSA (Hooke Laboratories, #DS-0162), emulsified with one volume of complete Freund’s adjuvant. Two subsequent boosts were performed at 21 days intervals, and an additional two 50 μg boosts after 16 months (see **Fig. 1A**). Production serum bleeds and bone marrow aspirates for V_H_H sequence libraries were obtained 10 days after the final boost. An HCAb fraction was prepared from serum bleeds through sequential binding on a Protein G column and then Protein A. Heparin was added to the HCAb fraction at a final concentration of 100 μg/mL and was incubated with magnetic streptavidin beads (Invitrogen, #11206D) coated with brominated 5’-biotin-(G:C)_30_ oligos (Integrated DNA Technologies). After thorough washing, the resin was resuspended in PBS and digested with IdeS (FabRICATOR, Genovis). Bound fragments were eluted in 1.1x LDS at 70 °C for 10 min, reduced and alkylated, and separated by SDS-PAGE. Bands corresponding V_H_H fragments (15-25 kDa) were excised from the gel, split into two portions and in-gel digested with either trypsin or chymotrypsin (Roche), and peptides were eluted and analyzed by LC-MS.

**Figure 1:**
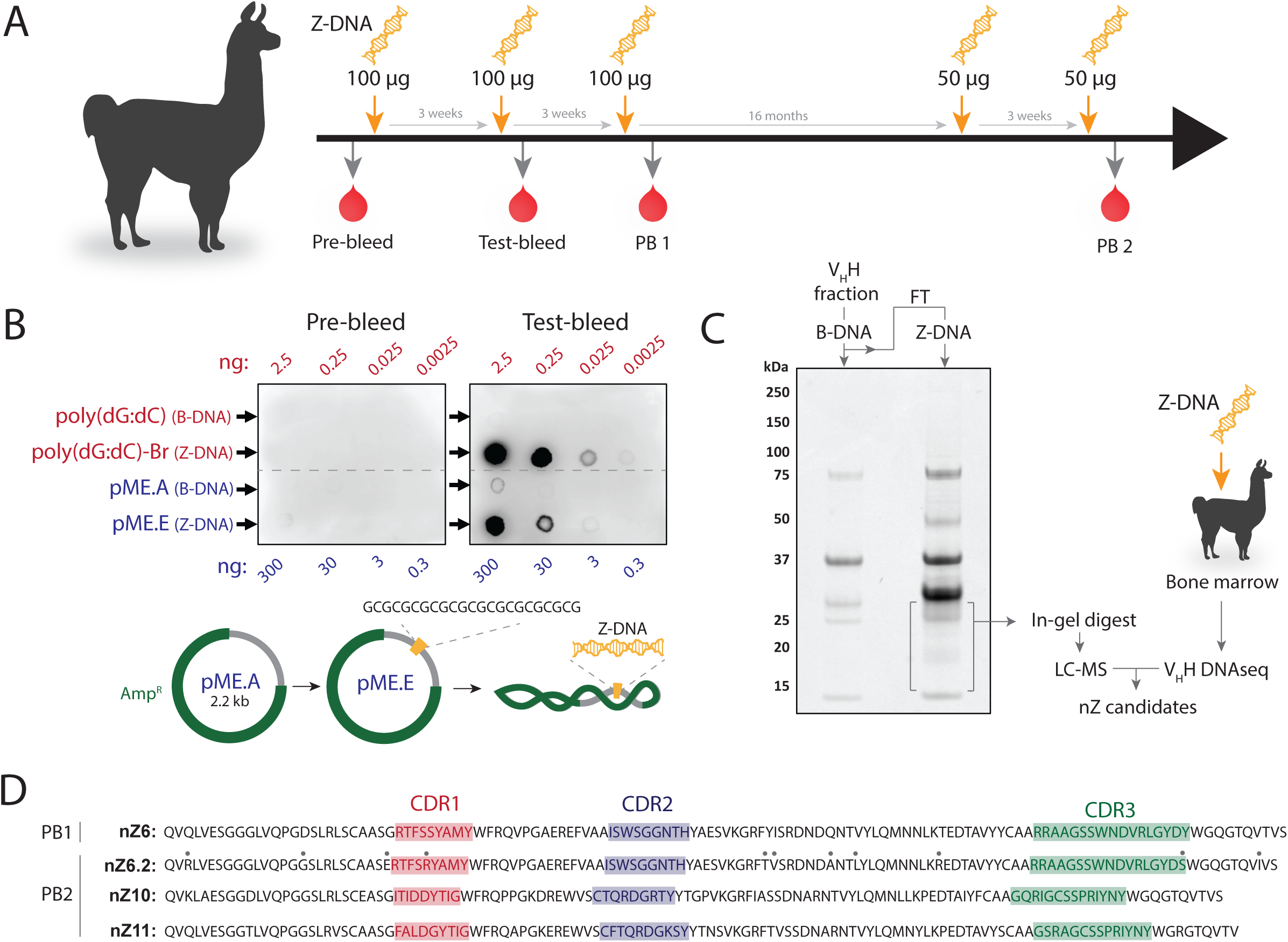
Isolation of Z-NA binding nanobodies. **A)** Overview of the immunization protocol used to generate Z-NA targeting nanobodies. A llama was immunized over three rounds with 100 µg brominated poly-(dG:dC) sequence (∽1.1kb average length), after which a firs production bleed (PB1) was collected. After 16 months, the same animal was exposed to another two injections of 50 µg of the same antigen, separated by three weeks, and followed by a second production bleed (PB2). **B)** Dot blots showing total Ig response in the animal before immunization (pre-bleed) and after the first two rounds of immunizations (test-bleed). The antigen used for immunization (brominated poly-(dG:dC); Z-DNA) was used as a positive control, while the non-brominated poly-(dG:dC) (B-DNA) was used as a negative control. A supercoiled plasmid with an insert of a 23 bp (dG:dC)-repeat sequence (pME.E) was used as a non-chemically modified positive substrate and the same plasmid without the insert (pME.A) was used as a negative substrate. The amount of DNA loaded on the blotting membrane is shown for both the linear substrates (red) and the plasmid substrates (blue). **C)** After pre-fractionation of the production bleed, to produce enriched HCAb, a magnetic resin coupled to non-brominated poly-(dG:dC) was first used to deplete specific and non-specific B-DNA binders from the HCAb fraction. The supernatant from this depletion was then combined with magnetic resin coated with brominated poly-(dG:dC), to enrich HCAb with specificity for Z-DNA. Bound HCAb was digested with IdeS, and remaining bound V_H_H fragments were eluted and run on SDS-PAGE. The bands in the V_H_H size region (∽15-25 kDa) eluted from the Z-DNA resin were excised and proceeded for peptide identification using LC-MS/MS. Nanobody candidates were identified by matching peptides against a total V_H_H DNA library sequenced from the immunized animal. **D)** The amino acid sequence of final four Z-binding nanobody candidates. One clone was isolated from PB1, while the last three were isolated from PB2. Clone nZ6.2 had an almost identical sequence with nZ6, but differed in 11 amino acid position (highlighted in gray circles above nZ6.2 sequence). The complementary determining regions are highlighted in red (CDR1), blue (CDR2), and green (CDR3).

### Cell culture

Immortalized N/TERT keratinocytes (N/TERT-2G) were used with permission from J. G. Rheinwald (Brigham and Women’s Hospital, Boston, MA, USA) and grown in KC serum-free medium (SFM) (Thermo Fisher Scientific, #17005-042) supplemented with bovine pituitary extract (30 μg/ml) (Thermo Fisher Scientific, #17005-042), epidermal growth factor (0.2 ng/ml) (Thermo Fisher Scientific, #17005-042), 0.1% (v/v) Amphotericin-B (Gibco, Thermo Fisher Scientific, #15290-018), 1% (v/v) Penicillin-Streptomycin (Gibco, Thermo Fisher Scientific, #15070-063) and calcium chloride (0.3 mM) (Gibco, Thermo Fisher Scientific, #50-9702). N/TERTs were used from passage 6 to passage 10. For IFN-α stimulation, cells were treated with IFN-α at a final concentration of 1000 U/ml (PBL assay science, 11100-1) in KC-SFM with added supplements for 16 hr.

### Immunoblotting

Dot blot experiments were performed on Nytran SPC nylon blotting membranes (Cytvia). The membrane was initially cut to size, loaded onto a vacuum-operated dot-blot apparatus, and each spot on the membrane was pre-equilibrated with 200 µL Tris buffer (10 mM, pH 8.0). Z-DNA substrate dilutions were prepared in the same Tris buffer at concentrations that would allow the desired substrate amount to be loaded onto the membrane in 100 µL. After the substrate solution had completely passed through the membrane, each spot was washed with 400 µL buffer. The membranes were then removed from the apparatus and blocked for 1 hr in 5% (w/v) non-fat milk in TBST (20 mM Tris, pH 7.6, 137 mM NaCl, 0.1% (v/v) Tween-20) at room temperature (RT). The blocking solution was the removed, and primary staining solutions containing either nanobody (nZ6, nZ6.2, nZ10, nZ11), hZɑ_ADAR1_-domain, or antibody (rabbit Z22, AbsoluteAntibody #Ab00783-23.0; mouse Z22, AbsoluteAntibody #Ab00783-2.0; sheep polyclonal IgG, Capralogics #P00008) were added. Primary detention reagents were used in a concentration of 1 μg/μl unless otherwise stated. Membranes were incubated with the primary staining solutions for 1 hr at RT. Membranes were then washed three times in TBST for 5 min before incubating with secondary staining solutions for 1 hr at RT (anti-rabbit IgG HRP, Cytiva #NA934, diluted 1:10.000; donkey anti-sheep IgG HRP, R&D Systems #HAF016, diluted 1:10.000; 6xHis monoclonal IgG HRP, Invitrogen #MA1-21315-HRP, diluted 1:5000). All primary and secondary detection reagents were diluted in 5% non-fat milk in TBST. Finally, membranes were washed four times with TBST for 5 min. An ImageQuant LAS-4000 system (GE Healthcare) was used for blot imaging on the standard setting with incremental image capture after the membrane had been treated with chemiluminescent HRP substrate (Millipore Sigma #WBLUF0100).

### Electrophoretic mobility shift assay

EMSAs were performed using fluorescently labeled, uniformly brominated on G nucleotides (positive control) and unbrominated (negative control) hairpin oligonucleotides. Sequence: 5’-FAM-ACG*CG*CG*CG*CG*TTTTCG*CG*CG*CG*CG*T-3’ (*indicates a G nucleotide +/-Br). Oligonucleotides (50 nM) were incubated with nZ6.2 at protein:oligonucleotide molar ratios of 20:1, 10:1, 5:1, and 2.5:1 (corresponding to 1 µM, 500 nM, 250 nM, and 125 nM nZ6.2, respectively) in 20 µL binding reactions containing 10 mM Tris (pH 7.5), 50 mM KCl, 3.5 mM DTT, 0.25% (v/v) Tween-20, and 0.05% (v/v) NP-40. Reactions were incubated for 1 hr at room temperature protected from light.

To assess the effects of oligonucleotide preconditioning, parallel reactions were performed using oligonucleotides preincubated for 1 hr in 32 mM Tris-HCl (pH 7.0), 330 mM KCl, and 1.8 mM cobalt hexammine at an oligonucleotide concentration of approximately 0.33 µM. Preconditioned oligonucleotides were added directly to binding reactions without buffer exchange, resulting in final concentrations of approximately 100 mM KCl and 0.27 mM cobalt hexammine. Protein binding was subsequently performed as described above. Following incubation, 5 µL Orange G/Ficoll loading buffer (15% (w/v) Ficoll 400) was added to each reaction. Samples were resolved on 7.5% native polyacrylamide gels (1.5 mm thickness) in 0.5× TBE buffer. Gels were prerun for approximately 30 min before sample loading, and electrophoresis was performed at approximately 120 V at room temperature protected from light. Fluorescent oligonucleotides were visualized using a fluorescence gel imaging system.

### Protein expression and purification

BL21-CodonPlus (Agilent) were transformed with pET28a+ vectors encoding 6-His tagged murine tandem ZBP1 Zα domains (Zɑ-Zɑ_Zbp1_; amino acids 1 to 153) as a wildtype construct or with point mutations in both Zα domains (mutZɑ-Zɑ_Zbp1_ = N46D, Y50A, N122D, Y126A (14, 48–50)). Single colonies were grown in 100 mL LB with antibiotics as preculture and were transferred to 2 L LB with antibiotics when the culture was turbid. The 2 L culture was incubated at 37°C 140rpm until an OD600 of 0.6 u was reached followed by a rapid cooling to 16 °C. IPTG was added to a final concentration of 1mM and the culture was further incubated overnight at 16°C. Bacteria were then pelleted and resuspended in NPI-10 (50 mM NaH2PO4, 300 mM NaCl, 10 mM Imidazole, 10 mM DTT) supplemented with 1 mM PMSF. Bacteria were then passed five times through an Avestin Emulsiflex C5 homogenizer for efficient lysis. The clarified lysate was incubated overnight at 4°C rotating with Ni-NTA resin (Qiagen) and applied to an empty chromatography column on the next day. The resin was washed once with NPI-10 containing 0.5% (v/v) Triton X-100, once with NPI-10 containing 0.25% (v/v) Triton X-100 and 1 M NaCl, twice with NPI-10 and eluted in NPI-250 (NPI buffer containing 250 mM Imidazole). Elution fractions were measured on a Nanodrop 8000 (Thermo Scientific) and analyzed by denaturing PAGE gel using a 10% solution of 2,2,2-Trichloroethanol (TCE, Sigma) for downstream protein staining and subsequent visualization on a UV gel imager. Elution fractions showing expression of the target size were pooled and concentrated with Amicon Ultra spin columns (10 kDa MWCO, Millipore) followed by size exclusion chromatography with DPBS (Gibco, Thermo Scientific) on an Äkta Pure system (Cytiva) with a Superdex 75 Increase 10/300 column (Cytiva).

BL21-Star-DE3 (Thermo) were transformed with pET28a+ vector encoding 6-His tagged human ADAR1 Zα domain (amino acids 133 to 209, a gift from Dr. Ky Lowenhaupt). Transformed cells were incubated for 1 hour in 750 µl SOC media, transferred to 20 mL LB with 50 µg/ml kanamycin pre-culture, and grown overnight to saturation at 37°C. The next day, 20 ml pre-culture was added to 1 L LB with kanamycin, incubated at 37°C 140rpm until an OD_600_ of 1 u was reached. IPTG was added to a final concentration of 200 µM and the culture was incubated four hours at 4 °C. Bacteria were then pelleted and flash frozen. Pellets were resuspended in 4 ml of lysis buffer per gram of cell pellet: 50 mM HEPES pH 8.0, 1 M NaCl, 20 mM Imidazole, 0.5 mM TCEP, and 0.1% (v/v) Tween-20 supplemented with 1 Pierce EDTA free protease inhibitor tablet per 50 mL of lysis buffer (Thermo). The cell suspension was passed twice through a French Press (Thermo), and the lysate was clarified by centrifugation, filtered through 0.2 µm, and purified over Ni-NTA resin (MCLAB). Resin was washed with 50 column volumes (CV) lysis buffer lacking protease inhibitors, 2 CV elution buffer with 20 mM imidazole, and then eluted with 4 CV elution buffer (20 mM HEPES pH 8.0, 150 mM NaCl, 300 mM Imidazole, 0.5 mM TCEP. The elution was loaded onto a 5 mL HiTrap heparin column (Cytiva) on an Äkta FPLC (buffer A: 20 mM HEPES pH 8.0, 150 mM NaCl, 0.5 mM TCEP and buffer B: 20 mM HEPES pH 8.0, 2M NaCl, 0.5 mM TCEP, 5% glycerol, 2 mM MgCl2). The protein eluted in two distinct peaks corresponding to 0.6 and 1.1 M NaCl, both of which were >99% pure Zα domain of ADAR1 as determined by SDS-PAGE. The tight binding fraction was concentrated to 13.4 mg/ml as determined by BSA standard using densitometry and used for all downstream experiments.

### Circular dichroism spectroscopy (CD)

All CD measurements were run using a Jasco J-715 instrument and a 0.1 cm quartz cuvette in 10 mM HEPES pH 7.4, 10 mM NaCl, 0.1 mM EDTA at room temperature (RT). DNA was mixed with tandem Zα domain constructs to a final concentration of 5 µM DNA and 20 µM protein and incubated for 30 min at RT. Measurements were performed between 220 nm and 320 nm with a data pitch of 1 nm, a scanning speed of 50 nm/min and a 1nm band with. CD spectra were generated from the average of three measurements. Control B-DNA spectrum was generated by measuring the d(CG)8 duplex mixed with PBS instead of protein and the Z-DNA spectrum was generated by incubating and measuring the d(CG)8 duplex in 4 M NaCl.

### Binding kinetic measurements by SPR

SPR experiments were performed on a Biacore 8K (Cytiva) instrument. Biotinylated (G:C)_30_ DNA oligos (+/-bromination) were immobilized at 100 ng/ml using the capture function set to 40 response units (RU) in PBS containing 0.05% (v/v) Tween-20 on a Series S SA sensor chip (Cytiva). Nanobodies were injected at 30 μl/min in multi-cycle kinetics experiments at concentrations between 0 and 640 nM, with an association time of 120 s, and a dissociation time of 240 s. Data were analyzed using Biacore software, and fitted using a two-step binding model. Sensor surfaces were regenerated between measurements with 0.1% (w/v) SDS. For epitope binning, pairs of binders were sequentially flowed using tandem dual injections according to the manufacturer’s guidelines. Proteins were injected at saturating concentrations with a flow rate of 10 µl/min. Contact time for the first protein was 120 sec, followed by 150 sec for the second protein, then a 30 sec dissociation. Response signal for the second binder was measured in a 10 sec window at the beginning of dissociation. Data were analyzed using the Biacore software epitope binning module.

### Immunoprecipitations

Tissue extracts and IPs were produced and conducted as previously described (51): the extraction solution was 20 mM HEPES pH 7.4, 300 mM NaCl, 1% (v/v) Triton X-100, supplemented with protease inhibitors; washing solutions were identical without protease inhibitors. IP samples used for spike-in experiments were prepared using 50 mg N2102Ep (**Fig. 3B-D**) or 50 mg N/TERT GFP-ZBP1 (**Fig. 3E**) cryomilled cell powder and 2.5 μl (slurry) Dynabeads M-270 Epoxy (ThermoFisher Scientific) conjugated with nZ6.2 or anti-GFP dimeric LaG16-LaG41 nanobody (46, 47). Clarified extracts were spiked with B- and Z-DNA (non-brominated or brominated poly(dG:dC)) and incubated for 30 min at 4 °C. In **Figure 3B-D**, extracts were spiked with 0, 100 or 500ng DNA, and in **Figure 3E**, extracts were spiked with 50 ng DNA. Affinity media was then added and the extracts were incubated at 4 °C for another hour and then washed three times with extraction solution and eluted with 1.1x NuPage LDS loading buffer at 70 °C. Eluted samples were either separated by SDS-PAGE and stained with Coomassie reagent (InstantBlue, Abcam #ab119211) in **Figure 3B**, analysed by LC-MS/MS in **Figure 3C-D**, or used for Western blotting of ZBP1 in **Figure 3E**.

### Mass spectrometry

For shotgun proteomic analyses of nZ6.2 immunoprecipitations, all proteins in the IP elutions were run on a NuPAGE 4-12% Bis-Tris gel (Invitrogen Novex). The gel was first pre-run empty for 3 min at 200V and then run with sample for 3 min at 200V. The gel was stained with InstantBlue Coomassie protein stain (Abcam #ab119211) for 1 hr and washed with Milli-Q water overnight. The stained gel region was sliced into small pieces, washed twice for 30 min at RT with mixing at 500 RPM (once with 30% and once with 50% (v/v) aqueous acetonitrile in 100 mM ammonium bicarbonate), and lastly with 100% acetonitrile for 5 min before drying (heat or vacuum). The proteins were reduced with 30 μL of 10 mM dithiothreitol (in 100 mM ammonium bicarbonate) for 30 min at 55 °C and then alkylated with 30 μL of 55 mM iodoacetamide (in 100 mM ammonium bicarbonate) for 30 min, in the dark, at RT. The gel pieces were then washed with 100% acetonitrile for 30 min while mixing (500 RPM) and dried before overnight digestion with 30 μL (12.5 ng/μl) trypsin. Digestion was carried out at 37 °C, overnight; the next day, the peptides were eluted from the gel pieces with 75% (v/v) acetonitrile, 5% (v/v) formic acid. The eluted fractions were dried under vacuum and resuspended in 0.1% (v/v) formic acid. Samples were analysed on an EASY-nLC 1200 system (Thermo Scientific) interfaced with an Orbitrap Exploris 480 (Thermo Scientific) using an EASY-Spray nano-electrospray ion source. The mass spectrometer was operated with a spray voltage of 1800 V, a capillary temperature of 305 °C. A precursor MS scan (m/z 350-1500, positive polarity) was acquired at a nominal resolution of 120K, followed by HCD-MS/MS at a nominal resolution of 30K. HCD was performed using a normalized collision energy of 24% and normalized AGC target was set to 100% with a maximum injection time of 116ms. MS raw files were analysed using FragPipe software (version 22.0) and searched against a human-specific proteomic database (UniProt, September 2024).

### Human subjects, ethics approval, and consent to participate

SLE patients and healthy controls were recruited through the University of Washington, Division of Rheumatology Biorepository. The study was approved by the University of Washington Institutional Review Board and informed written consent was obtained from all participants according to the Declaration of Helsinki. The study was approved by the University of Washington Institutional Review Board. Informed written consent was obtained from all participants according to the Declaration of Helsinki.

### Neutrophil isolation

Neutrophils were isolated from freshly drawn blood by gradient centrifugation on PolymorphPrep (CosmoBio, USA) according to the manufacturer’s instructions. The cells were washed and suspended in DMEM high glucose. By flow cytometry using anti-CD66b, neutrophil purity was >95%.

### UV irradiation of N/TERTs

N/TERTs were seeded at a density of 250,000 cells per well on uncoated glass coverslips (22 mm by 22 mm; Thermo Fisher Scientific, #3406) in six-well plates and allowed to adhere for 24 hr. Cells were left untreated (mock) or primed with IFN-α for 16 hr. On the day of UVB irradiation, medium was removed and replaced with 1 ml of prewarmed 1x PBS per well, and cells were irradiated with a dose of 50 mJ/cm² using the UV-2 ultraviolet irradiation system (Tyler Research; emission at 310 nm via cascade-phosphor ultraviolet generators), or left untreated. Immediately after irradiation, PBS was removed and replaced with 2 ml of fresh prewarmed medium per well.

### Immunofluorescence microscopy

Freshly isolated neutrophils were cultured for 30 min at 37°C on 96-well glass bottom plates (Cellvis #P96-1-N) pre-coated with poly-L-lysine (Millipore-Sigma # P4707). The adherent neutrophils were fixed with 4% paraformaldehyde solution for 20 min, followed by two washes with phosphate-buffered saline (PBS). Cells were permeabilized with 0.1% Triton X-100 in PBS for 10 min followed by three washes with PBS. To prevent nonspecific binding, neutrophils were blocked using 10% normal mouse and/or goat serum in BlockAid solution with anti-human FcgRII antibody overnight at 4°C. The cells were incubated with nanobody at 10 μg/mL for 2h and subsequently stained with goat anti-mouse IgG secondary Ab conjugated with Alexa Fluor 488 (green) or Alexa Fluor 647 (red). Negative controls consisted of cells stained with secondary antibodies alone. Nuclei were stained with either DAPI or Hoechst33342. Images were captured at 40 X, 60 X, or 100 X magnification using a Keyence BZ-X800 Fluorescence microscope and analyzed using BZ-X800 Analyzer software.

For Z-DNA staining of N/TERTs, cells were fixed 3 hr after UVB exposure with freshly prepared 4% formaldehyde solution (Thermo Scientific, 3215658) (at RT for 15 min; immunofluorescence was performed on the same day as the experiment. After fixation, cells were washed and permeabilized with PBS + 0.1% Triton X-100 (Gibco, Thermo Fisher Scientific, #10010023) (Sigma, T8787) (wash buffer). Cells were then blocked with 5% Bovine Serum Albumin (Fischer Bioreagents, BP1600-100) and 10% normal rabbit serum (Jackson ImmunoResearch, 177613) in washing buffer (blocking buffer) for 30 min at RT. Primary antibody cocktails were prepared in blocking buffer and incubated on each coverslip for 1 hr at RT. This cocktail includes nZ6.2 nanobody (1:500), and TOMM22 (1:200) (Abcam, AB57523). Wells were washed three times with wash buffer and incubated with secondary antibodies diluted in block buffer for 30 min at RT. The secondary antibody cocktail includes DAPI (1:1000) (Invitrogen, 62248), Camelid VHH antibody (1:500) (GenScript, A01862-200), and Alexa Fluor 647 goat anti-mouse (Invitrogen, A21241). After three additional washes with wash buffer, coverslips were mounted on glass slides using ProLong Glass Antifade Mountant (Invitrogen, P36980). The brightness and contrast of the nZ6.2 channel were adjusted for visualization purposes.

## Results

### Isolation of Z-NA binding nanobodies

To generate nanobodies that specifically recognize Z-DNA, we used a similar approach as was used for generating the original Z22 antibody (27). In the immunization protocol (**Figure 1A**), a brominated (dG:dC)n repeat sequence of about 1.1kb was initially injected into a llama, followed by two subsequent booster immunizations. After the first two immunizations, a test bleed was performed to assay the initial immune response in the animal. The test bleed serum reactivity was compared to pre-bleed serum from the same animal, obtained prior to the initial immunization (**Figure 1B**). Reactivity was assessed by dot blot using the brominated DNA immunogen, cognate unbrominated DNA control, and negatively super-coiled plasmids with (pME.E, Z-DNA) or without (pME.A, B-DNA) a 23bp (dG:dC) insert (52). In both cases, test-bleed reactivity was very strong for positive control substrates while showing little to no reactivity against negative control substrates. Pre-bleed serum did not display any reactivity toward any of the tested substrates. We then employed a proven method for selecting high-affinity nanobody candidates that involves liquid chromatography-coupled tandem mass spectrometry (LC-MS/MS) based peptide mapping to a V_H_H DNA sequence library (46, 47). A heavy-chain antibody (HCAb) serum fraction was first incubated with a B-DNA-coated affinity resin ((dG:dC)_60_), and the flow-through fraction was then incubated with a separate resin coated in Z-DNA (brominated (dG:dC)_60_). Fc domains were released with an IdeS digestion, and V_H_H domains from each of the bound fractions eluted with SDS and run on SDS-PAGE (**Figure 1C**). All Z-DNA-enriched bands in the V_H_H size region were in-gel digested and peptides were mapped using LC-MS/MS, using a V_H_H DNA sequence library generated from the immunized animal; four nanobody candidates were selected for further validation. After recombinant production and purification, the candidates were screened based on their ability to bind test substrates by dot blotting. Here, only one of the candidates - nZ6 - bound to both the brominated poly(dG:dC) substrate and the pME.E plasmid (**Figure 1D & S1**). In order to increase the success rate and generate additional Z-DNA binders, the same animal was injected with two additional booster shots of brominated DNA (**Figure 1**). From the second production bleed, 22 candidates were selected for further screening, and of these, three nanobodies displayed selective binding to the test substrates in the dot blot screen (**Figure 1D & S2**). One of the nanobodies - nZ6.2 - has a very high sequence similarity to nZ6 (91.2%). We therefore contend that it is very likely to be a rediscovery of an affinity matured HCAb derived from the nZ6 nanobody. The two other nanobodies - nZ10 and nZ11 - also have high sequence similarity in their CDR3 loops which suggests that they may interact with identical epitopes on the Z-DNA structure.

### *In vitro* binding characteristics to synthetic substrates

Next, we used dot blot assays to characterize the nanobodies’ binding attributes, probing immobilized poly(dG:dC) substrates (1 - 0.04 ng) and the supercoiled pME plasmids (200 - 8 ng). **Figure 2A** compares the nanobodies and the hZɑ_ADAR1_-domain (**Figure S3**), as a positive control, at a fixed concentration (1 µg/mL). As expected from their sequence similarities, nZ6 and nZ6.2 appear to bind similarly to the two substrates although nZ6.2 produces a stronger signal overall, which is further evidence of affinity maturation. Likewise, nZ10 and nZ11 produce very similar binding patterns, which is congruent with their high CDR3 similarity. Unlike nZ6 and nZ6.2, they both appear to bind more favorably to the brominated substrate. This difference in the binding patterns between nZ6/nZ6.2 and nZ10/nZ11 could be explained by different Z-DNA epitopes. It is also possible that the nZ10/nZ11 pair is more reliant on a structural feature that is more prevalent in the brominated poly(dG:dC) sequence, or upon the Br substituent itself. None of the four nanobodies exhibited detectable binding to control B-DNA substrates at the tested concentration in the dot blot assay.

**Figure 2:**
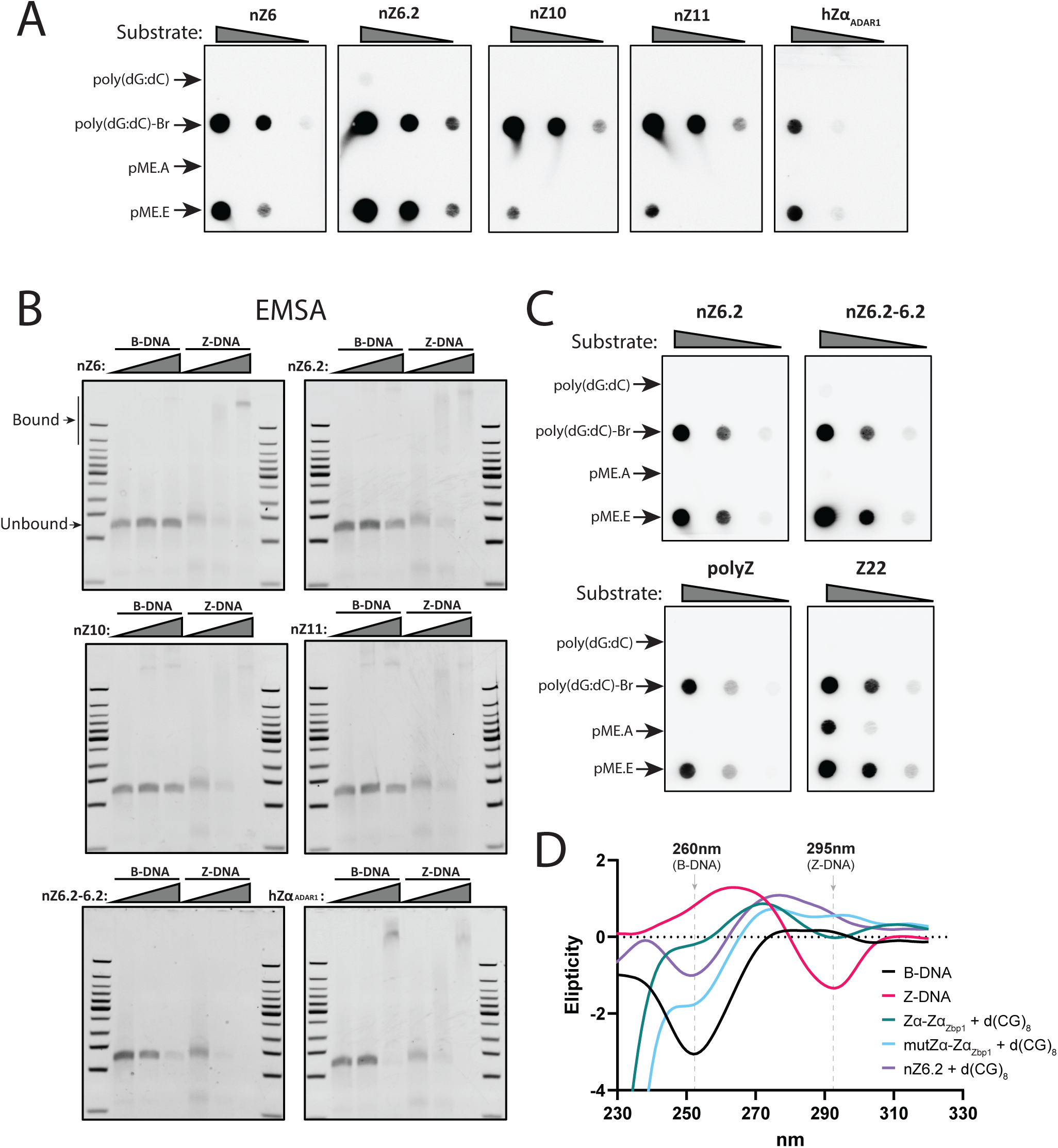
In vitro binding characteristics of Z-binding nanobodies. **A)** Dot blots using linear poly-(dG:dC) substrate with (Z-DNA) and without (B-DNA) bromination and supercoiled plasmid with (Z-DNA, pME.E) and without (B-DNA, pME.A) a 23 bp (dG:dC)-repeat insert. Linear substrate quantities: 1, 0.2 and 0.04 ng; plasmid substrate quantities: 200, 40 and 8 ng. All nanobodies and hZɑ_ADAR1_ (‘Zɑ’) were applied a 1 µg/mL concentration. **B)** Electrophoretic mobility shift assay on 65 nM brominated (Z-DNA) and non-brominated (B-DNA) (dG:dC)_30_ probes incubated with 0, 500, and 2000 nM nanobody or hZɑ_ADAR1_. **C)** Dot blot stainings using linear poly-(dG:dC) substrate with (Z-DNA) and without (B-DNA) bromination and supercoiled plasmid with (Z-DNA, pME.E) and without (B-DNA, pME.A) a 23 bp (dG:dC)-repeat insert. Linear substrate quantities: 1, 0.2 and 0.04 ng; plasmid substrate quantities: 200, 40 and 8 ng. Monomeric nZ6.2, dimeric nZ6.2-6.2, polyclonal anti-Z-DNA antibody (polyZ), and monoclonal Z22 antibody were used at a concentration of 1 µg/mL. **D)** Circular dichroism spectroscopy measurements of a (dC:dG)_8_ DNA duplex mixed with the indicated tandem hZɑ_ZBP1_-domain (WT and binding-deficient mutant (mut Zɑ)) or nZ6.2 nanobody. B-DNA control in black corresponds to (dC:dG)_8_ incubated in PBS without protein and the Z-DNA control in red corresponds to (dC:dG)_8_ in 4M NaCl. The B-DNA mark indicates the characteristic valley of the B-DNA spectrum at ∼260nm while the Z-DNA mark indicates the Z-DNA characteristic valley at ∼295nm.

Binding between the nanobodies and Z-DNA was also tested using electrophoretic mobility shift (EMSA) assays. Here, a fixed concentration (65 nM) of control or brominated (dG:dC)_30_ probe was incubated with different concentrations (0, 0.5, or 2 µM) of nanobody or hZɑ_ADAR1_-domain and electrophoretic shift was observed using non-denaturing polyacrylamide gel electrophoresis and SYBR Gold nucleic acids staining (**Figure 2B**). All four nanobodies produced a concentration dependent shift in the Z-DNA probe accompanied by the concomitant disappearance of the unbound probe signal. The nZ10 and nZ11 nanobodies also show a fainter band shift of the B-DNA probe. In an effort to make a Z-NA binder with higher affinity, the nZ6.2 clone was homodimerized using a 4xG_4_S-linker. When this nZ6.2-6.2 dimeric nanobody was tested using the EMSA assay, we saw a complete shift of the Z-DNA probe already at the 0.5 µM nanobody concentration, although no higher migrating band could be observed: the dimeric nature of the nanobody and the repeating epitope structure of the substrate give rise to substrate crosslinking possibilities, producing higher-order complexes that may not migrate into the polyacrylamide gel matrix. Combining 2 µM nZ6.2-6.2 with B-DNA yielded similar results. As has been shown in several other studies (21–23, 53, 54), the Zɑ domain of human ADAR1 (hZɑ_ADAR1_-domain) is capable of binding both B- and Z-DNA probes at sufficiently high concentration. While the probe shift is more apparent for Z-DNA, the hZɑ_ADAR1_-domain also completely shifts the B-DNA probe at the highest concentration. Comparable EMSA results were also obtained in separate experiments combining nZ6.2 and a hairpin DNA substrate with (Z-DNA) or without (B-DNA) uniform bromination of G residues (**Figure S4**). These results also revealed: (1) the hairpin Z-DNA shifts are consistent with multiple nanobodies per DNA substrate at higher nanobody:substrate ratios (**Figure S4A)**; and (2) the nanobody can bind and shift a B-DNA substrate, if the substrate has been pre-incubated at sufficiently high salt concentration to facilitate the B→Z shift, e.g., aided by cobalt hexamine chloride. (6, 55–58) (**Figure S4B)**.

For further characterization we decided to focus primarily on the nZ6.2 clone because this nanobody demonstrated excellent detection capabilities without any apparent binding differential between brominated poly(dG:dC) and non-brominated pME.E Z-DNA substrates, and without any discernible cross-reactivity with B-DNA at normal working concentrations. We next compared the Z-DNA detection sensitivity of both the nZ6.2 monomer and dimer against the Z22 monoclonal antibody and a sheep polyclonal antibody. Using the Z-DNA dot blot assay (**Figures 2C and S5**), we see that all reagents bound the Z-DNA substrates in a comparable manner. However, unlike the other tested reagents, the Z22 antibody also bound the supercoiled control plasmid without the GC-repeat insert; an observation that we encountered in several experiments using the Z22 antibody.

An important feature to be aware of, for any Z-NA binding reagent, is whether it binds to the Z-NA structure after the B→Z transition (selective for Z-NA), or if it can also actively induce the B→Z transition in a concentration dependent manner, as has been described for Zɑ-domains (21–25). To determine if the nanobody possessed B→Z conversion capabilities, we used CD spectroscopy to measure the absorbance of a B-DNA probe in the presence of nZ6.2 (**Figure 2D & S6**). B-DNA spectral traces will exhibit a characteristic valley at ∼260 nm, while Z-DNA traces exhibit a valley at ∼295 nm. When we incubated a B-DNA probe with a purified tandem Zbp1 Zɑ domain (Zɑ-Zɑ_Zbp1_; see Methods) we observed that the resulting spectral trace exhibited a valley at ∼295 nm; incubating B-DNA with a purified mutant construct that is deficient for B→Z conversion (mutZɑ-Zɑ_Zbp1_; see Methods), mitigated the formation of this characteristic valley. In contrast, when the B-DNA probe was incubated with the nZ6.2 nanobody, the resulting curve more closely matched the characteristics of the B-DNA control, supporting the conclusion that the nZ6.2 nanobody, unlike the Zɑ-domain, does not readily induce the B→Z transition.

To further compare our cohort of nanobodies to Z22 and hZɑ_ADAR1_, we measured their binding kinetics by surface plasmon resonance (SPR), using immobilized DNA with and without bromination. Along with the nanobodies, we also measured the binding affinity of Z22 and the hZɑ_ADAR1_-domain (**Table 1, Figure S7A**). We found that nZ6 and nZ6.2 had higher K_D_s (respectively, ∼70 nM and ∼50 nM) than nZ10 and nZ11 (both ∼7 nM). However, it is notable that the dot blotting results for nZ10 and nZ11 demonstrated superior binding of these reagents to brominated Z-DNA substrates, compared with the non-modified, Z-DNA-containing plasmid substrate; nZ6.2 showed excellent binding to both the brominated DNA and the Z-DNA plasmid (**Fig. 2A**). When homodimerized, the affinity of nZ6.2-6.2 (K_D_ ∼20 pM) increased several orders of magnitude compared to the monomeric version. For hZɑ_ADAR1_, we measure a binding affinity of ∼6 nM, which is comparable to what has been described previously (13). To assay the different binding features of these reagents, we checked their ability to simultaneously interact with the brominated DNA substrate, in pairwise comparisons (potentially indicating that they bind at different locations). We found that each reagent blocked the subsequent reagent from binding; this was true of the nanobodies, Z22, and the hZɑ_ADAR1_-domain (**Fig. S7B**).

#### Functional validation of nZ6.2 binding to Z-DNA in complex biological samples

Having validated nZ6.2’s Z-DNA binding capabilities *in vitro,* using purified components in low complexity settings, we next assayed the nanobody’s ability to selectively retrieve spiked-in Z-DNA (brominated poly(dG:dC) substrate) from a cell lysate, thereby testing its potential as an immunoprecipitation (IP) reagent (**Figure 3A**). For this, aliquots of a clarified lysate from N2102Ep cells were spiked with either B-DNA or Z-DNA and incubated at 4 °C for 30 mins - to provide an opportunity for proteins from the complex mixture to bind the DNA (whether specifically or non-specifically), potentially blocking nZ6.2 from binding in an IP context. After this, nZ6.2-conjugated magnetic beads were also added to the mixture and incubated together at 4 °C for 1 hr; because nZ6.2 selectively binds Z-DNA, we expected to pull down proteins that bind to Z-DNA, only in the samples spiked with Z-DNA. As a readout, proteins eluted from all the samples were visualized by SDS-PAGE, followed by Coomassie blue staining (**Figure 3B**). The Z-DNA-spiked, affinity-enriched fractions retrieved a range of different proteins, whose abundances corresponded with the spike-in amount; contrastingly, the B-DNA-spiked fractions did not retrieve proteins at abundances detectable with Coomassie blue staining. We profiled the non-spiked and Z-DNA-spiked IP fractions by LC-MS/MS and observed a dramatic enrichment (>250x) of ADAR1 (**Figure 3C**). Inspecting the peptide coverage of ADAR1 in the spiked samples, we found several peptides matching the ADAR1 Zɑ-domain, supporting specific enrichment of the ADAR1 p150 isoform to the recovered spiked-in Z-DNA (**Figure 3D & Supp. Table S1**). We did not observe ZBP1-derived peptides. To verify whether nZ6.2 can enrich ZBP1 through Z-DNA, we used an N/TERT1 keratinocyte cell line engineered to constitutively overexpress a GFP-tagged *ZBP1* gene fusion. As with the N2102Ep lysates, clarified N/TERT1 cell lysates were spiked with either B- or Z-DNA and nZ6.2 was used for IP, with the expectation that ZBP1 should co-IP from Z-DNA-spiked but not B-DNA-spiked or non-spiked samples. Western blotting demonstrated that GFP-ZBP1 was recovered in the Z-DNA-spiked lysate, but not from the B-DNA-spiked or non-spiked controls (**Figure 3E**). In addition, the recovery of ZBP1 in the IP was sensitive to the NaCl concentration in the extraction solution, co-IPing with Z-DNA at 150 mM but not 250 mM NaCl.

**Figure 3:**
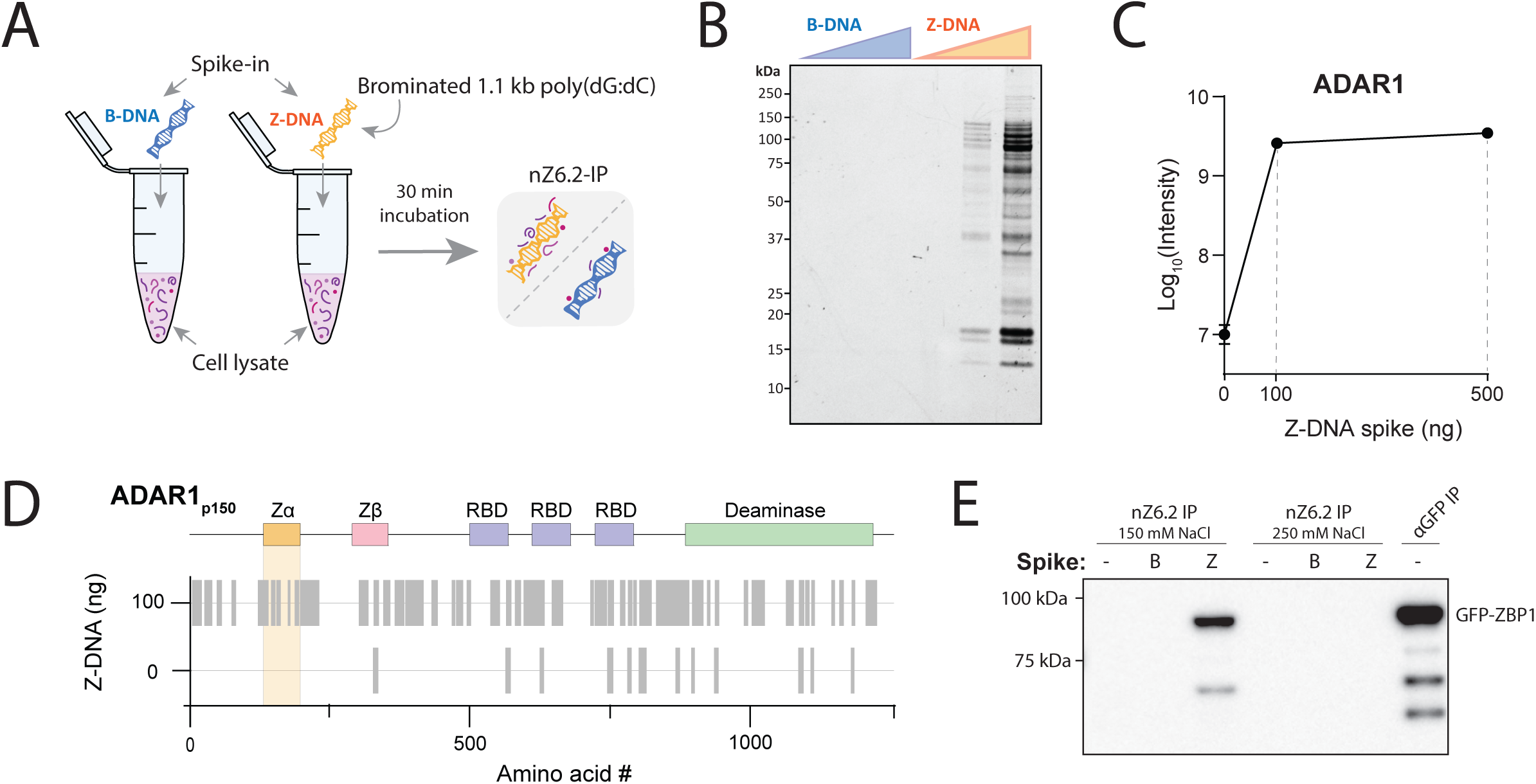
Z-NA immunoprecipitation using nZ6.2. **A)** Total cell extracts were spiked and incubated with brominated (Z-DNA) or non-brominated (B-DNA) poly-(dG:dC) substrate (∽1.1kb). Substrates with bound proteins were recovered by immunoprecipitation using magnetic beads conjugated with nZ6.2 nanobody. **B)** Coomassie-stained (Abcam InstantBlue^®^) SDS-polyacrylamide gel of eluted proteins from N2102Ep extracts spiked with 0, 100, or 500 ng substrate (B-or Z-DNA). **C)** ADAR1 protein intensities measured by LC-MS in nZ6.2 IPs from N2102Ep cell extracts spiked with different amounts of Z-DNA (n=2). **D)** ADAR1 protein coverage map, showing LC-MS/MS identified ADAR1 peptides (gray bars) in nZ6.2 IPs from Z-DNA-spiked (0 and 100ng) N2102Ep extracts. Peptides are mapped against a schematic drawing of the protein domains in ADAR1. The Z-NA binding Zɑ-domain is highlighted in yellow. **E)** Western blot probed for ZBP1. The probed samples are nZ6.2 IP pullouts of B-or Z-DNA-spiked extracts from N/TERTs overexpressing GFP-ZBP1. Protein extracts and nZ6.2 IPs were done at 150 and 250 mM NaCl. An anti-GFP IP was performed at 250 mM NaCl as a positive control.

#### Immunofluorescence staining comparisons between nZ6.2 and Z22

Next, we tested the binding of nZ6.2 to endogenous Z-NA substrates. Recently (manuscript submitted), we have observed that endogenous Z-NAs are able to trigger interferon production in neutrophils isolated from patients with systemic lupus erythematous (SLE): accordingly, probing SLE neutrophils with nZ6.2 revealed prominent cytoplasmic signal that substantially overlapped with ZBP1 signals (**Figure 4A**). Neutrophils isolated from healthy controls did not show any positive staining from nZ6.2 (**Figure S8**). When the Z-NA positive SLE neutrophils were probed with both nZ6.2 and Z22 reagents, they produced directly adjacent signals and exhibited only minor overlap (**Figure 4B**). Testing clone nZ11 on SLE neutrophils revealed a comparable pattern to that observed with clone nZ6.2 but at weaker overall signal strength (**Figure S9**), in agreement with these reagents’ respective performance binding unmodified Z-DNA (compare poly(dG:dC)-BR with pME.E, **Figure 2A**). A majority of SLE patients experience cutaneous IFN-triggering photosensitivity, most likely caused by ultraviolet B (UVB) light-linked mitochondrial damage in keratinocytes. Recent findings show that mitochondrial damage coincides with the co-accumulation of Z-DNA and ZBP1 (17). Based on this direct evidence of a stimulus triggered Z-DNA accumulation in keratinocytes, we tested whether the nZ6.2 clone could detect this induction of Z-form species. N/TERT keratinocytes were pre-treated for 16 hr with IFNɑ, before being dosed with UVB light. Both treated and untreated cells were stained with nZ6.2 and TOMM22 as a mitochondrial marker. The treated N/TERT cells show a marked increase in diffuse punctate nZ6.2 staining throughout the cytoplasm when compared to untreated control (**Figure 4C**).

**Figure 4:**
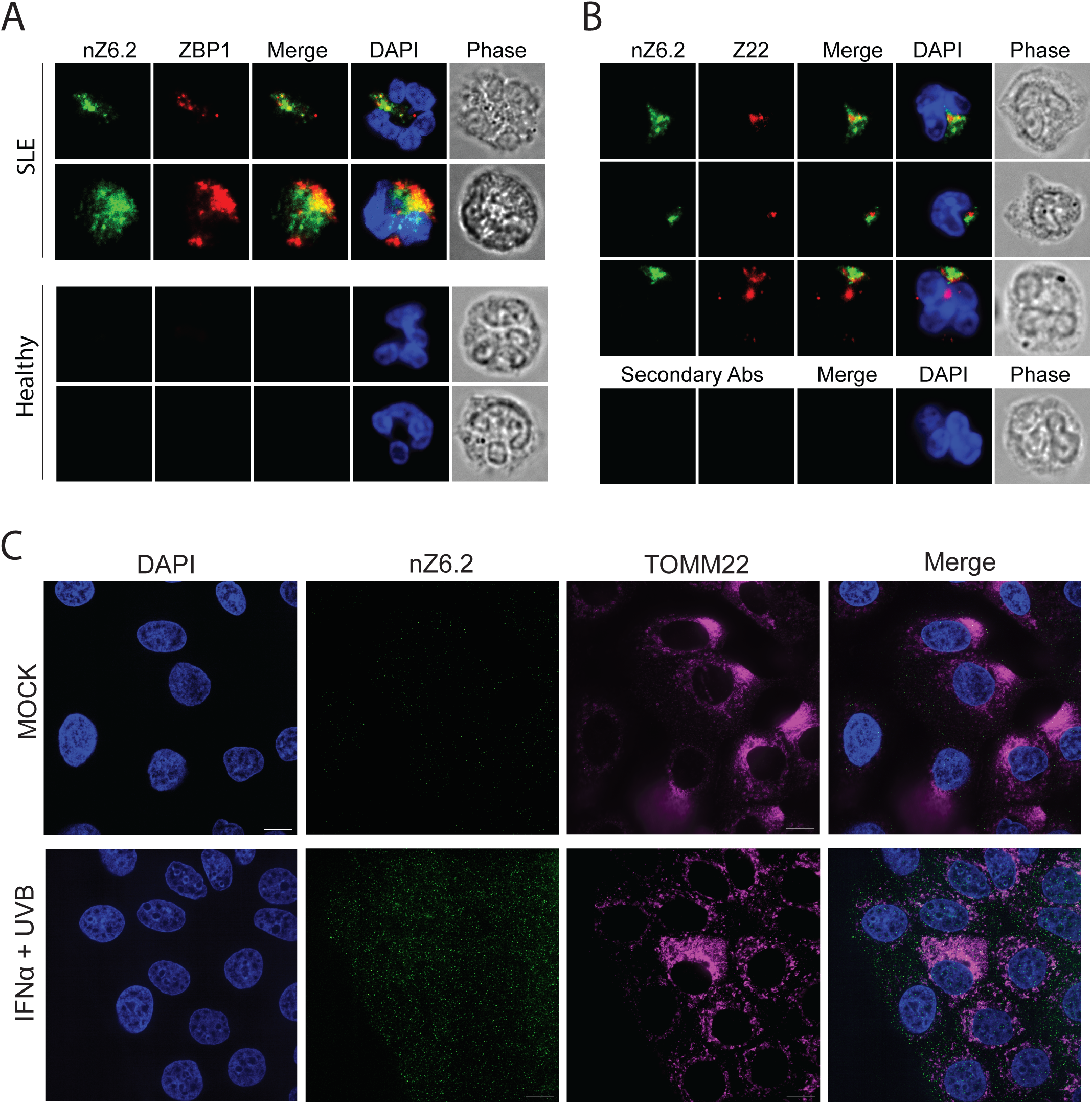
Immunofluorescence microscopy of SLE patient neutrophils and N/TERT cells using nZ6.2. **A)** Three representative SLE neutrophils stained with nZ6.2 (green), anti-ZBP1 (red), and DAPI for DNA (blue). The third image in each row is a merge of green and red, the fourth includes DAPI, and the fifth is a phase contrast of the same cell. The bottom row is a SLE neutrophil stained with secondary antibodies alone. **B)** Same experiment with Z22 (red) instead of anti-ZBP1. These images represent more than 50 neutrophils each from 5 independent SLE patients. **C)** Representative images of N/TERT cells treated with 1000 U/mL IFN-α and 50 mJ/cm^2^ UVB irradiation or untreated mock, stained with DAPI, nZb6.2 nanobody (iFLuor488), and TOMM22 antibody (AlexaFluor 647). Scale bar = 10 μm.

## Discussion

In this study, we generated and characterized a new cohort of recombinant llama-derived nanobodies that recognize Z-DNA. Four clones (designated nZ6, nZ6.2, nZ10, and nZ11) were identified following immunization with brominated poly(dG:dC), affinity enrichment, and mass spectrometry-guided clone discovery (**Fig. 1**). Among these, nZ6.2 emerged as the most broadly useful reagent, combining strong affinity for both brominated and non-brominated Z-DNA substrates with minimal detectable B-DNA cross-reactivity (**Figs. 2** & **S5; Table 1**). Importantly, unlike canonical Zα domains – which recognize Z-NA and lower the energetic barrier for B/A→Z transitions – nZ6.2 did not promote B→Z conformational conversion, indicating preferential recognition of pre-existing Z-DNA (**Figs. 2D & S6**). Notably, a Z22-derived chimeric Fab has also recently been shown to directly induce a concentration-dependent B→Z transition in CG-repeat DNA *in vitro* (35); to our knowledge this has not been directly assayed for the original full-length murine Z22 IgG2b. In contrast, some single-molecule and biochemical studies have posited an alternative mechanism involving conformational selection and stabilization of transiently formed Z-DNA (59, 60); Z-DNA recognition and B-to-Z conversion can be experimentally separable activities, as demonstrated for different Zα domains (25) and the anti-Z-DNA clone D11antibody (31). Reagents that perturb the structural equilibrium of their targets can complicate interpretation of localization and capture experiments. The apparent inability of nZ6.2 to induce Z-DNA therefore suggests that it may provide a less perturbative means of interrogating endogenous Z-NA biology, particularly in biochemical enrichment and imaging applications. Z22 exhibited an impressive ∼50 pM K_D_ between Z22 and brominated poly(dG:dC) Z-DNA; prior studies also measured relatively high affinities (15.6 nM K_D_ (_61_) and 9.1 nM K_D_ (_13_), using monovalent Z22 Fab rather than bivalent Z22 IgG). However, the Z22 antibody also bound to the pME.A B-DNA control plasmid (compare pME.A control plasmid to pME.E Z-DNA plasmid for nZ6.2 and Z22, **Fig. 2C**), and SPR measurements demonstrated low but discernable Z22 signals for B-DNA binding (**Figure S7).** The observed Z22:B-DNA signals, measured by maximum binding (Rmax, in response units (RU)), are only ∼2% of those exhibited by Zα, perhaps representing fractionally induced Z-form upon Z22 binding; but the measured affinity, if these signals are interpreted, is significant (106 pM, or ∼½ the affinity seen for brominated poly(dG;dC)). These results suggest that Z22 IgG has a higher Z-DNA affinity than nZ6.2 but lower selectivity for only pre-formed Z-DNA. Of note, our nanobodies all exhibited affinities within the same order of magnitude of Z22 Fab (compare **Table 1** to the prior studies (13, 61)).

The ability of immobilized nZ6.2 to selectively enrich Z-DNA together with known Z-NA-binding proteins, ADAR1 and ZBP1 (**Fig. 3**), demonstrates that the nanobody functions effectively as an affinity capture reagent, retrieving Z-form nucleoprotein assemblies (‘ZNPs’) from complex biological mixtures; this nominates nZ6.2 for future affinity ‘omic studies targeting endogenous Z-NAs (e.g., by IP-MS and IP-seq). Corroborating the IP-based result, and providing a disease-relevant, physiological context, we observed substantially overlapping staining patterns when contrasting nZ6.2-vs. ⍺-ZBP1-stained SLE-patient-derived neutrophils (**Fig. 4A**). In the keratinocyte SLE skin model, the primary target of cytosolic ZBP1 binding is posited to be oxidized (8-OH-dG) Z-DNA of mitochondrial origin (17); therefore nZ6.2’s ability to detect endogenous Z-NA structures in keratinocytes further supports its utility for cellular imaging in SLE disease models (**Fig. 4C**). Although nZ6.2 and Z22 produced proximal staining patterns in SLE derived neutrophils, their only partial overlap (**Fig. 4B)** may reflect distinct epitopes or different Z/B-DNA selectivities. They may also exhibit distinct modes of target accessibility as Z-NAs are decorated with endogenous proteins within the cellular milieu; each reagent may cope with these decorations differently, and/or have different degrees of selectivity for e.g., Z-DNA vs. Z-RNA. Yet, this minimal overlap additionally underscores the value of having independent reagents for studying these nucleic acid structures in cellular contexts.

Taken together, our research has established a collection of renewable, robust recombinant binders that complement existing Z-NA detection reagents and expand the experimental toolkit available for studying Z-NAs. nZ6.2 and its higher affinity dimer are favorably positioned for future use as ‘omics affinity reagents and for targeted investigations of the spatiotemporal dynamics of Z-form nucleic acids during innate immune signaling and inflammatory responses. Until now, investigations of Z-NA biology have relied heavily upon a small number of monoclonal antibodies, particularly Z22, which has ultimately become the de facto standard and the only reagent that is widely distributed commercially. Dependence primarily upon a single reagent limits opportunities for orthogonal validation and restricts rigorous research to applications compatible with the sole reagent. In contrast to traditional IgGs, recombinant nanobodies offer several practical advantages, including defined sequence, renewable production, ease of engineering, and compatibility with intracellular expression and fusion-protein reporter constructs (44). We note that the paratope of Z22 has been defined (31, 35) and a Z22 scFv has been reported (61, 62). While nanobodies still hold an edge over scFvs in many applications, both reagents could potentially be engineered as fusion proteins with comparable reporters and used essentially identically in common applications, improving head-to-head comparisons and the discernment of their relative advantages.

Interest in Z-form nucleic acids has increased substantially following the recognition of ZBP1 and ADAR1 as central regulators of innate immunity and inflammatory disease. Continued progress will require collections of orthogonal molecular tools that allow Z-NA capture and detection across many distinct experimental frameworks. Equally important are tools that can discriminate Z-RNA and Z-DNA from one another, without inducing B/A→Z transitions. The recombinant nanobodies described here expand the repertoire of available Z-NA binders and provide versatile, renewable reagents for studying Z-NA in biochemistry and molecular cell biology.

### Limitations of this research and future directions

The biochemical and biophysical characterization presented here focused primarily on synthetic DNA substrates; additional studies will be required to define binding to Z-RNA structures that are increasingly implicated in innate immunity. For example, while the bound cytosolic targets revealed in the SLE skin model are oxidized mtDNAs (17), the bound cytosolic targets in SLE neutrophils are less certain (and are speculated to potentially be DNA:RNA hybrids produced by dysregulated LINE-1 reverse transcriptase activity (T. Mustelin, personal communication; manuscript submitted). Hence, although the staining patterns we observed in immunofluorescence imaging experiments were consistent with endogenous Z-NA detection, the precise molecular identities of the cellular structures recognized has not always been firmly established; this is largely true of the many different cellular contexts explored across the corpus of cytosolic Z-NA research where the targets were probed using Z22.

Differences observed between distinct nanobody’s binding attributes suggest that multiple Z-NA epitopes or conformational states may exist, warranting future structural characterization; indeed, prior studies have revealed such distinctions between Z-NA binders (27, 30, 34, 63). However, our own attempts to bin our nanobody reagents, Z22, and hZɑ_ADAR1_ by their binding activities, using competitive SPR assays, revealed that most were at least partially interfering with one another on a 60 bp brominated poly(dG:dC) substrate (**Figure S7B**). An interesting case is presented by the distinction between Z-NA binding and the binding of B→Z junctions (i.e., the locations of structural transition between the forms); Z22 has been reported to have a 100-fold stronger affinity for B→Z junctions than continuous Z-DNA stretches (64). With this in mind: we observed that nZ6/6.2 nanobody clones bind brominated 1.1 kb poly(dG:dC) substrate with a comparable end-point intensity as the Z-DNA-forming pME.E plasmid (see **Fig. 2A** [dot blot]), yet the nZ10/11 nanobody clones show a clear preference for brominated poly(dG:dC); in SPR nZ10/11 nanobody clones exhibited several-fold higher affinity for 60 bp brominated poly(dG:dC) compared with nZ6/6.2. The differences may be explained by a preference of nZ10/11 for brominated vs. unmodified Z-NA, or they could also plausibly be explained by a preference of nZ10/11 for B→Z junctions: the pME.E plasmid likely has just two prevalent B→Z junctions (flanking the dG:dC repeat sequence, at the boundaries with the rest of the plasmid), whereas the brominated substrates could have many such transitions depending on the degree of bromination (which is not strictly defined in these substrates) and the substrate length. Our dot blots did not reveal detectable Z22 binding to unmodified 1.1 kb poly(dG:dC) DNA (B-DNA), while clear binding to brominated poly(dG:dC) Z-DNA was observed. At the same time, Z22 exhibited binding to pME.A (B-DNA; **Fig. 2C**).

On the whole, there is much room for improvement in the development of high affinity and highly selective ⍺-Z-NA reagents that can target and discriminate different Z-NA forms, as well as the substructures that may exist within them. Because nanobodies are genetically encoded single-domain binders, they are highly engineerable; it is foreseeable that we may be able to recover e.g., Z-RNA-vs. Z-DNA-selective clones by mutagenesis and screening of our current nanobody sequence library, raised against brominated poly(dG:dC) DNA (65). We also intend to administer Z-RNA and Z-DNA:RNA hybrid boosters to llamas that were previously responsive to Z-DNA. Moving ahead, applications we intend to explore and further develop with these reagents, their derivatives, and future clones we may produce, include engineered multivalent constructs with altered avidity (comparable process to the nZ6.2 dimer), nanobody-fluorescent protein fusions for live-cell imaging (a.k.a., intrabodies or chromobodies (66, 67)), *in situ* proximity-labeling and IP-MS for true endogenous (rather than spike-in based) interactomics. Such applications should facilitate mechanistic studies of Z-NA formation, turnover, and signaling during transposon expression, viral infection, interferon responses, and autoimmune diseases.

## Supporting information

LEGENDS for Sup Figs

Fig S1_nanobody candidates 1

Fig S2_nanobody candidates 2

Fig S3_Z-alpha domain

Fig S4_EMSA

Fig S5_Dot blots

Fig S7_SPR

Fig S8_neutrophils_additional

Fig S9_neutrophils_nZ11

Table S1_ADAR1_MS

## Data Availability

The data used in this article are available in the article and in its online supplementary materials. Raw mass spectrometry and imaging data files will be shared on reasonable request to the corresponding author.

## Supplementary Files

**Figure S1**: Nanobody candidate screening dot blots from PB1

**Figure S2** :Nanobody candidate screening dot blots from PB2

**Figure S3**: Coomassie gel on purified hZɑ_ADAR1_-domain

**Figure S4**: Additional nZ6.2 EMSA gels using fluorescent oligonucleotide probes

**Figure S5**: Additional dot blot comparisons using nZ6.2, nZ6.2-6.2, Z22, polyclonal anti-Z-DNA,

**Figure S6**: Additional data for CD spectroscopy on nZ6.2 and Zɑ-Zɑ_Zbp1_

**Figure S7**: SPR binding curves and binning table

**Figure S8:** Additional IF stainings of healthy and SLE neutrophils using nZ6.2

**Figure S8:** IF co-stainings of SLE neutrophils using nZ11 and Z22

**Table S1:** LC-MS data on ADAR1 from nZ6.2 IPs on Z-DNA spiked cell extracts.

## Author Contributions

**Conceptualization**: M Nielsen, J LaCava; **Data curation**: M Nielsen, P Fridy, L Wachsmuth, H Jiang, T Mustelin; **Formal analysis**: M Nielsen, P Fridy; **Funding acquisition**: JM Kahlenberg, T Mustelin, J LaCava; **Investigation**: M Nielsen, P Fridy, J Da Silva, M Pourshamsi, L Wachsmuth, J Kittel, H Jiang, BD Miller; **Methodology**: M Nielsen, L Wachsmuth, J LaCava; **Project administration**: M Nielsen, J LaCava; **Resources**: MS Taylor, BT Chait, M Pasparakis, JM Kahlenberg, MP Rout; **Supervision**: MS Taylor, M Pasparakis, JM Kahlenberg, T Mustelin, J LaCava; **Validation**: M Nielsen, H Jiang, J LaCava; **Visualization**: M Nielsen, P Fridy, J LaCava; **Writing – original draft**: M Nielsen, J LaCava; **Writing – review & editing**: M Nielsen, P Fridy, BD Miller, MS Taylor, JM Kahlenberg, T Mustelin, MP Rout, J LaCava

## Acknowledgements

We thank Mrs. Kelly R. Molloy for advice and assistance in nanobody candidate selection and Dr. Ky Lowenhaupt who generously shared historical reagents that aided this work, including the discontinued Amersham Biosciences poly(dG-dC)·poly(dG-dC) product (#27-7910-03) and the plasmid encoding recombinant hZɑ_ADAR1_. We thank the patients and healthy volunteers who donated blood for this study.

