## Supplementary material for "New llama-derived nanobodies enable selective detection and capture of Z-DNA": LEGENDS for Sup Figs

### Supplementary figure legends

**Supplementary Figure S1**: Dot blots for PB1 nanobody candidate pre-screening. Membranes were loaded with 1 ng linear poly-(dG:dC) substrate without (B-DNA) or with (Z-DNA) bromination, and 200 ng supercoiled plasmid without (B-DNA, pME.A) or with (Z-DNA, pME.E) a 23 bp (dG:dC)-repeat insert. Blots were incubated directly with filtered culture media (of unquantified nanobody concentration) harvested five days post-transfection of Expi-HEK293T cells with constructs encoding nZb1, 2, 3, 4, 6, 8, and 9. Controls were incubated with polyclonal anti-Z (positive) or media from mock-transfected cells (negative).

**Supplementary Figure S2**: Dot blots for PB2 nanobody candidate pre-screening. Membranes were loaded with 1 ng linear poly-(dG:dC) substrate without (B-DNA) or with (Z-DNA) bromination, and 200 ng supercoiled plasmid without (B-DNA, pME.A) or with (Z-DNA, pME.E) a 23 bp (dG:dC)-repeat insert. Blots were incubated directly with filtered culture media (of unquantified nanobody concentration) harvested five days post-transfection of Expi-HEK293T cells with constructs encoding nZb6.2, nZb10–26, and nZb28–30. Controls were incubated with polyclonal anti-Z.

**Supplementary Figure S3**: Purity assessment of recombinantly produced human ADAR1 Zα domain (hZ$\alpha_{ADAR1}$). Coomassie-stained SDS-PAGE gel showing heparin column elution fractions of purified hZ$\alpha_{ADAR1}$ alongside BSA standards (0.5, 2, 8, and 12 µg) and protein molecular weight ladders.

**Supplementary Figure S4**: Electrophoretic mobility shift assays (EMSA) of nZ6.2 binding to Z-DNA versus B-DNA. Fluorescently labeled 10 bp GC-repeat hairpin probes, either brominated on guanine residues (Z-DNA oligo) or unmodified (B-DNA oligo), were incubated with increasing molar ratios of nZ6.2 nanobody (2:1, 5:1, 10:1, and 20:1; NA indicates probe-only control) under standard buffer conditions **(A)** or in the presence of 330 mM KCl and 1.8 mM $[Co(NH_3)_6]Cl_3$ **(B)**. Arrows indicate free probes as well as 1:1 and 2:1 nZ6.2:oligonucleotide complexes.

**Supplementary Figure S5**: Dot blot comparison of Z-DNA binding reagents across three independent replicates (Replicates #1–3). Membranes were spotted with serial dilutions of linear 1.1 kb poly-(dG:dC) control (B-DNA; 1, 0.2, and 0.04 ng, row A), brominated 1.1 kb poly-(dG:dC) (Z-DNA; 1, 0.2, and 0.04 ng, row B), control plasmid pME.A (B-DNA; 200, 40, and 8 ng, row C), and 23 bp GC-insert plasmid pME.E (Z-DNA; 200, 40, and 8 ng, row D). Blots were probed with monovalent nZ6.2 (1, 0.2, and 0.1 µg/mL), bivalent nZ6.2-6.2 (1 and 0.2 µg/mL), polyclonal anti-Z (polyZ; 1 µg/mL), rabbit Z22 (1 µg/mL), or mouse Z22 (1 µg/mL).

**Supplementary Figure S6**: Circular dichroism (CD) spectroscopy of nZ6.2 and Zbp1 Zα domains with a poly-(dG:dC) probe (CG8). **(A)** Two independent experimental replicates (Rep#1 and Rep#2) showing CD spectra (230–320 nm) of B-DNA and Z-DNA controls alongside CG8 incubated with nZ6.2, wild-type mouse Zbp1 tandem Zα domain (Zα-Z$\alpha_{Zbp1}$), or binding-deficient mutant (mutZα-Z$\alpha_{Zbp1}$), as well as protein-only controls. **(B–E)** Individual subsets of curves replotted from Rep#1 in **(A)** to facilitate direct visual comparison of protein-only controls **(B)**, all protein + CG8 incubations **(C)**, Zα-$Z\alpha_{Zbp1}$ and mutZα-$Z\alpha_{Zbp1}$ conditions **(D)**, and nZ6.2 conditions **(E)**.

**Supplementary Figure S7**: Surface plasmon resonance (SPR) binding kinetics and epitope binning of Z-DNA binders. **(A)** Multi-cycle SPR sensorgrams showing dose-dependent binding of nanobodies (nZb6, nZb6.2, bivalent 2xnZb6.2, nZb10, and nZb11), Z22 antibody, and human ADAR1 Zα domain ($Z\alpha$) to immobilized brominated poly-(dG:dC) (Z-DNA) versus non-brominated poly-(dG:dC) (B-DNA) across the indicated concentration series. **(B)** Pairwise epitope binning matrix illustrating competitive co-binding on Z-DNA between sequential injections of Binder 1 and Binder 2, colored from blocking (red) to non-blocking (blue).

**Supplementary Figure S8**: Immunofluorescence (IF) microscopy of Z-DNA in human neutrophils using nZ6.2. Representative images of neutrophils isolated from systemic lupus erythematosus (SLE) patients **(A)** and healthy donors **(B)** co-stained with nZ6.2 nanobody (green), Z22 antibody (red), and DAPI for DNA (blue), alongside merged and phase-contrast channels. Secondary-antibody-only controls are shown for both groups.

**Supplementary Figure S9**: Immunofluorescence (IF) microscopy of Z-DNA in SLE patient neutrophils using nZ11. Representative images of neutrophils from SLE patients co-stained with nZ11 nanobody (green), Z22 antibody (red), and DAPI for DNA (blue), alongside merged and phase-contrast channels.
