## Supplementary figures and images for "New llama-derived nanobodies enable selective detection and capture of Z-DNA"

### Fig S1_nanobody candidates 1

#### Supplementary Figure S1

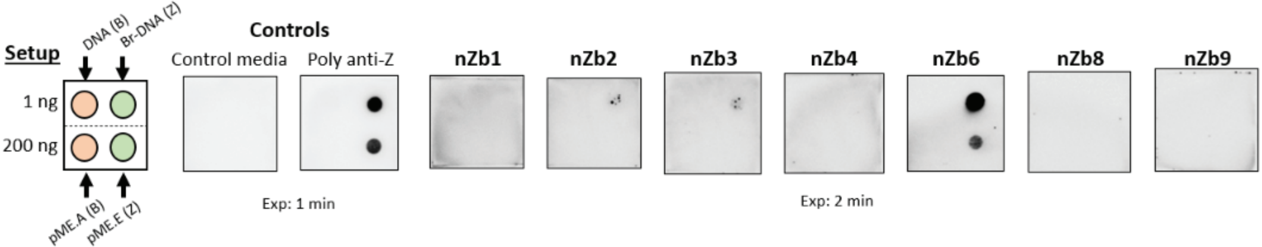

### Fig S2_nanobody candidates 2

#### Supplementary Figure S2

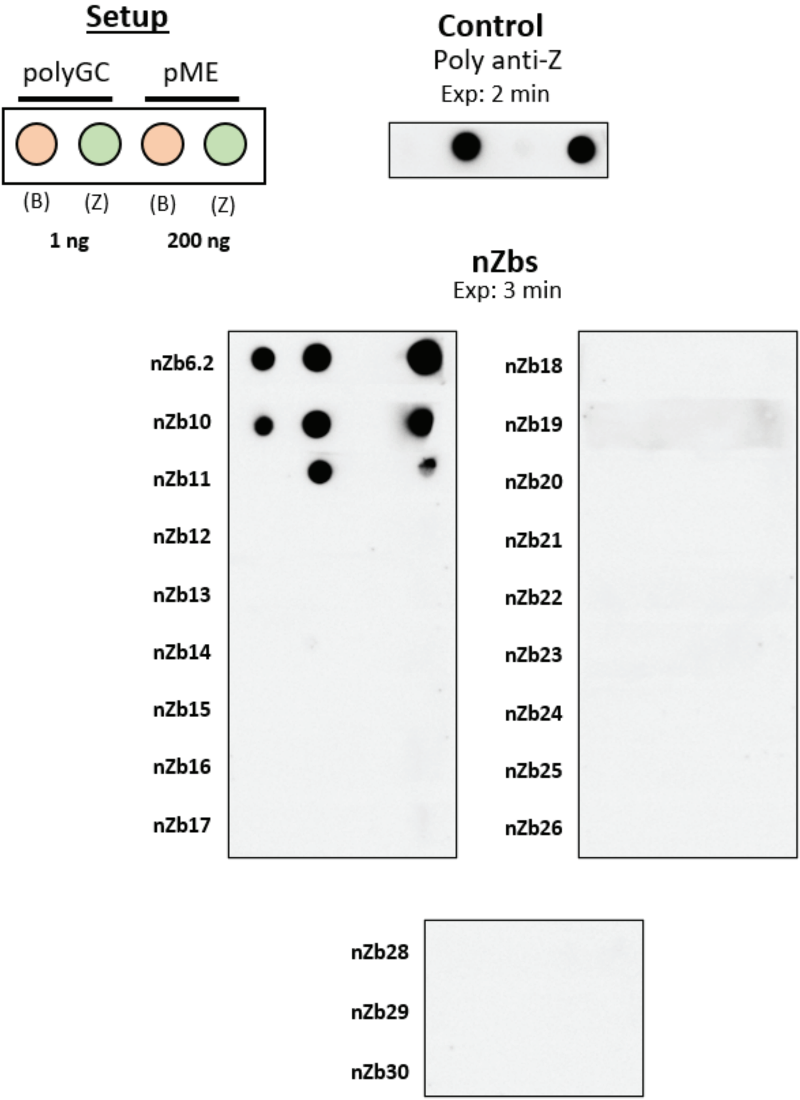

### Fig S3_Z-alpha domain

#### Supplementary Figure S3

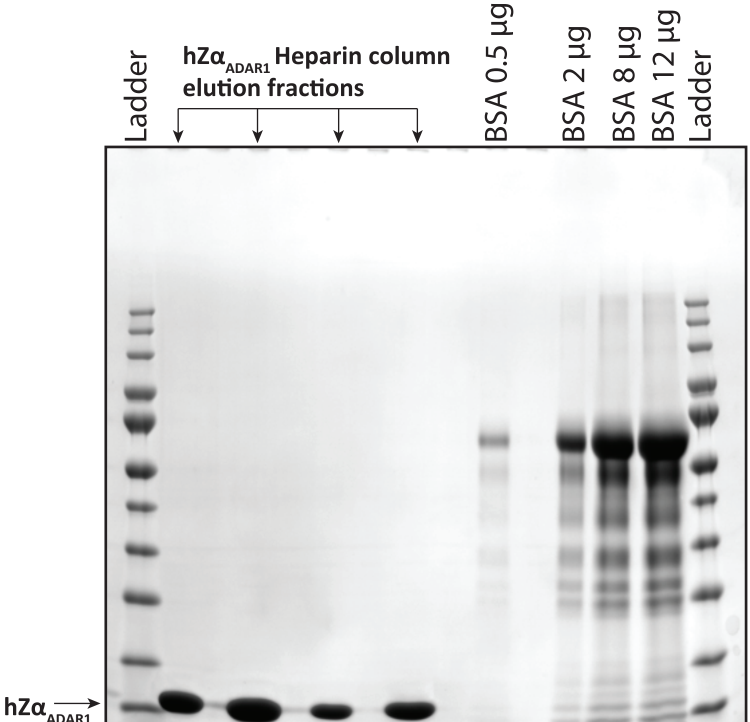

### Fig S4_EMSA

#### Supplementary Figure S4

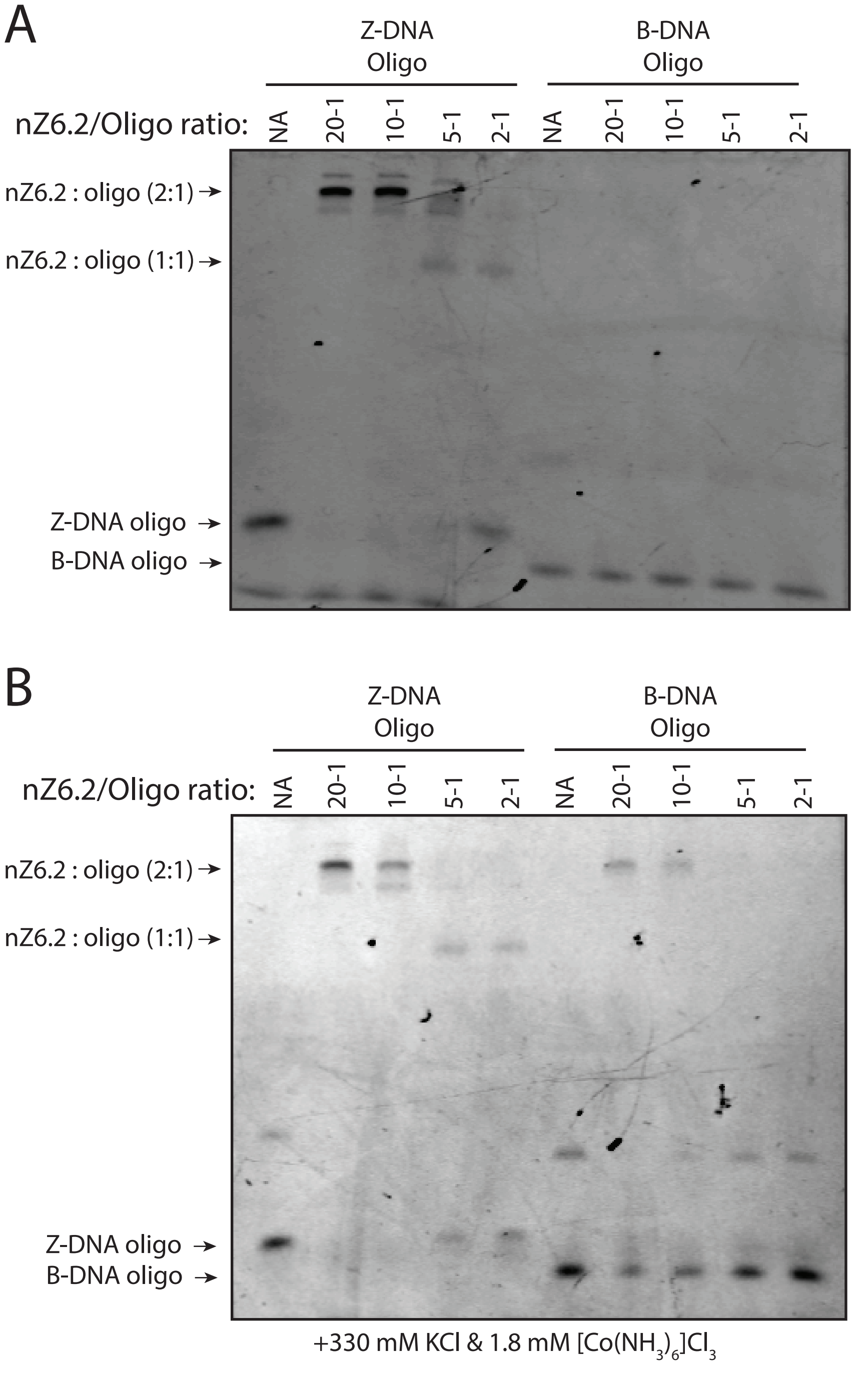

### Fig S6_CD

#### Supplementary Figure S6

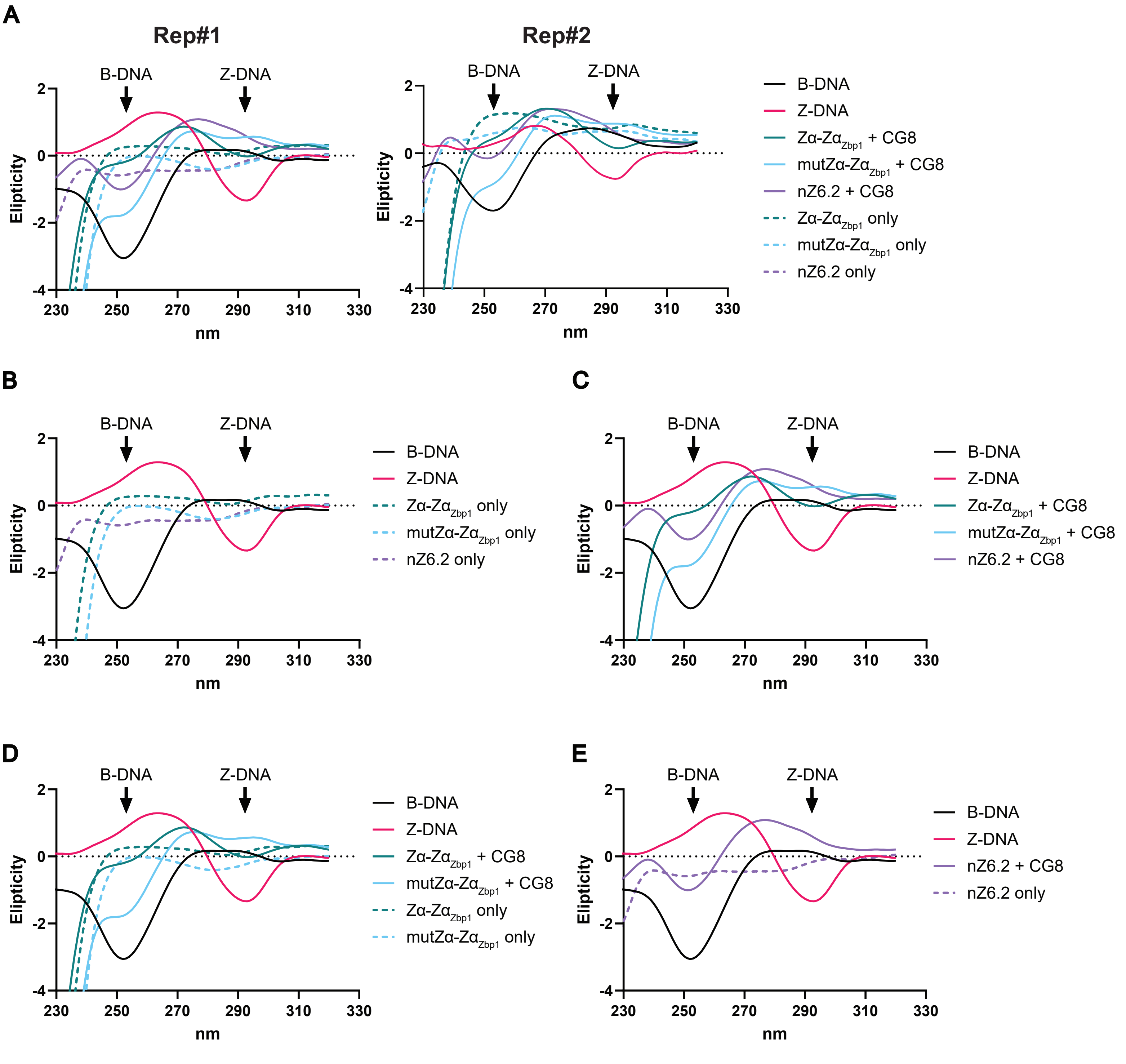

### Fig S7_SPR

#### Supplementary Figure S7

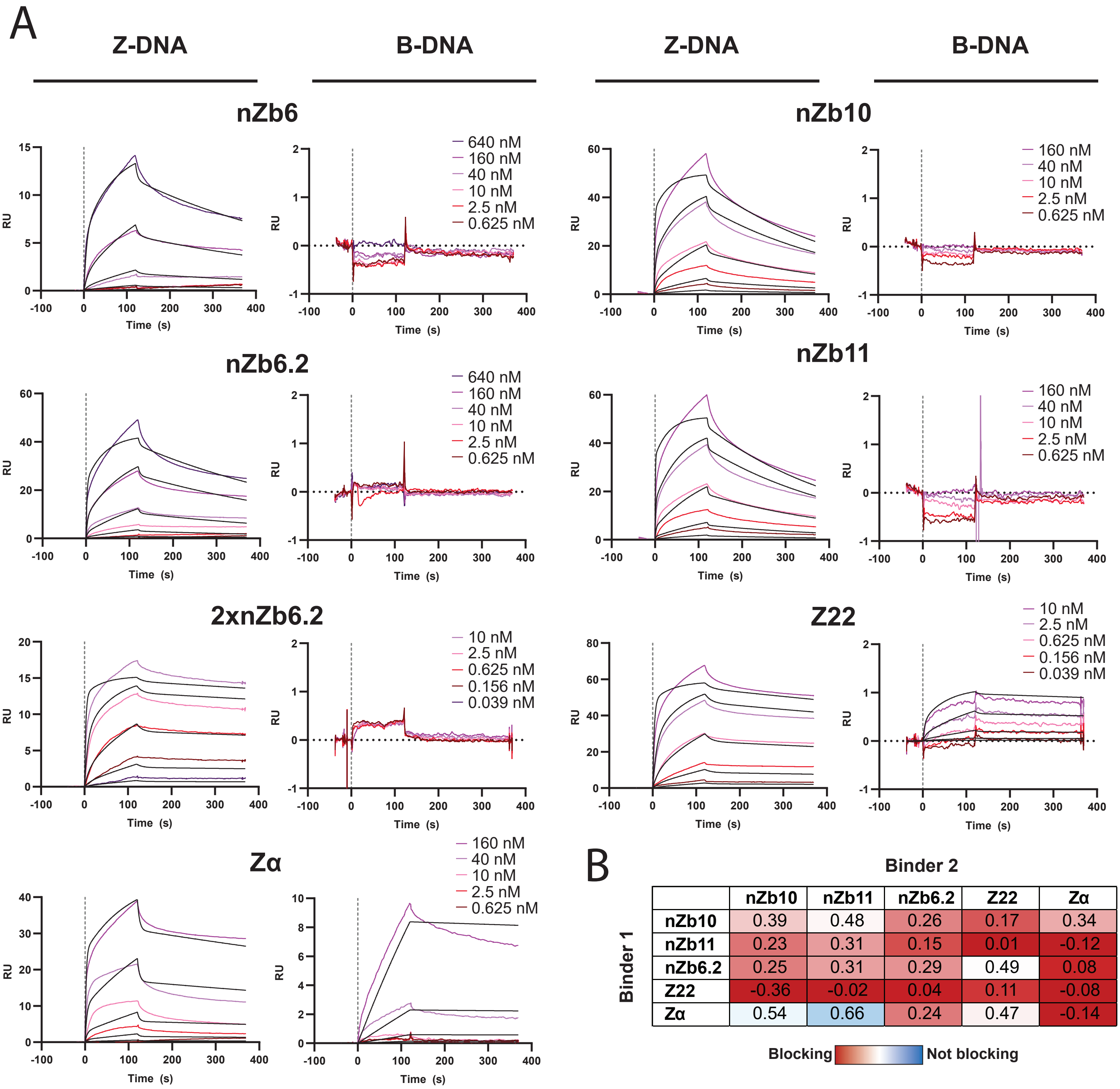


B

Binder 2

| Binder 1 | nZb10 | nZb11 | nZb6.2 | Z22 | Zα |
|---|---|---|---|---|---|
| nZb10 | 0.39 | 0.48 | 0.26 | 0.17 | 0.34 |
| nZb11 | 0.23 | 0.31 | 0.15 | 0.01 | -0.12 |
| nZb6.2 | 0.25 | 0.31 | 0.29 | 0.49 | 0.08 |
| Z22 | -0.36 | -0.02 | 0.04 | 0.11 | -0.08 |
| Zα | 0.54 | 0.66 | 0.24 | 0.47 | -0.14 |

Blocking — Not blocking

### Fig S8_neutrophils_additional

#### Supplementary Figure S8

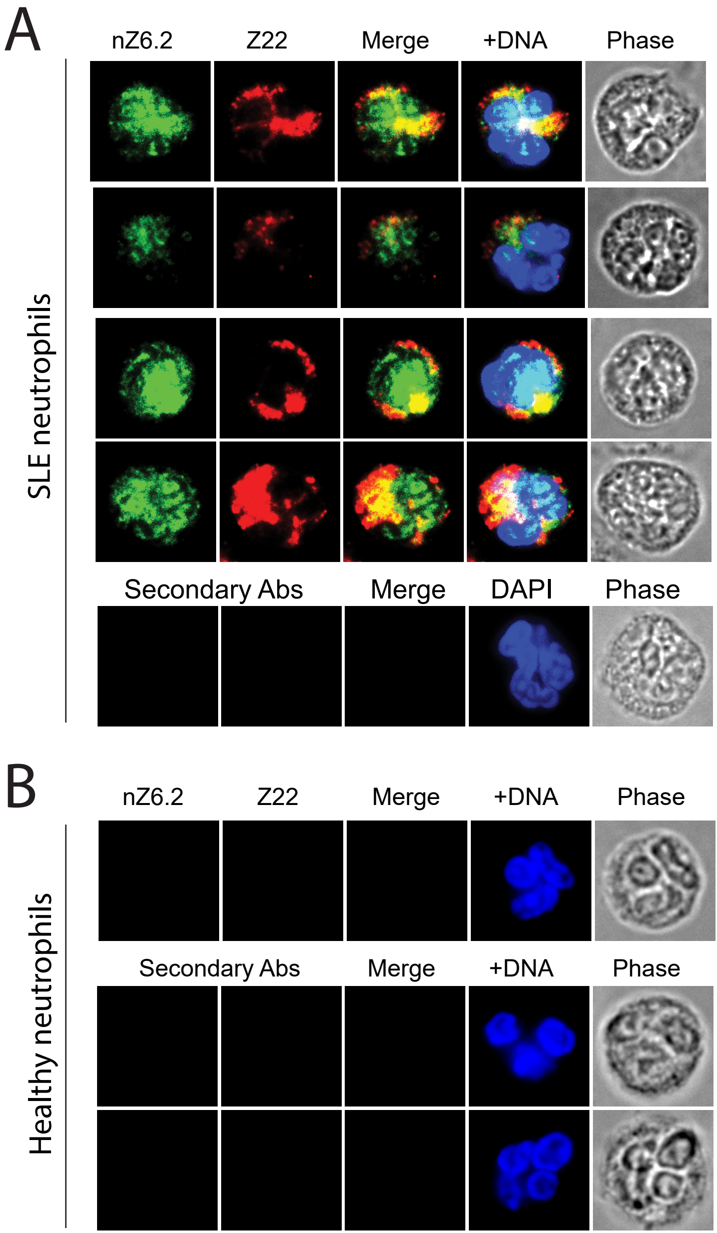

### Fig S9_neutrophils_nZ11

#### Supplementary Figure S9

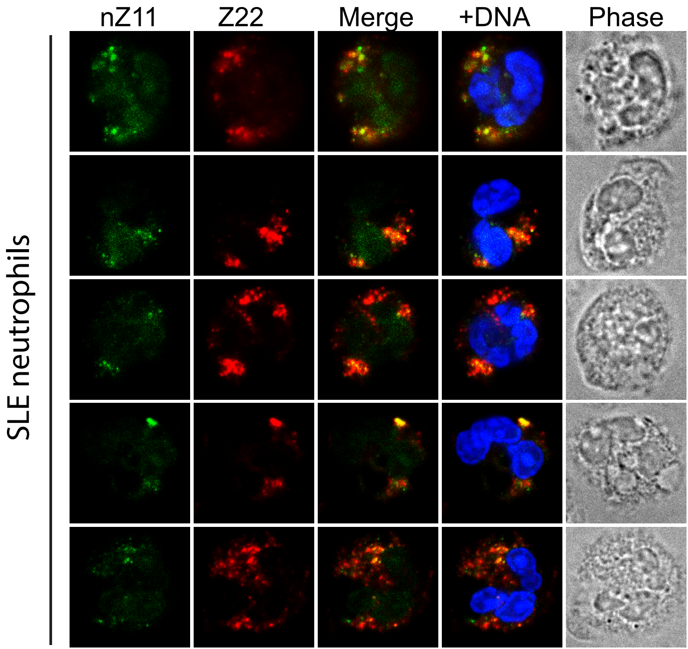
