## Supplementary material for "New llama-derived nanobodies enable selective detection and capture of Z-DNA": Fig S5_Dot blots

### Supplementary Figure S5

A: B-DNA 1.1 kb poly(dG:dC) control (1, 0.2, 0.04 ng)
B: Z-DNA 1.1 kb poly(dG:dC) brominated (1, 0.2, 0.04 ng)
C: B-DNA pME.A control plasmid (200, 40, 8 ng)
D: Z-DNA pME.E 23bp GC insert plasmid (200, 40, 8 ng)

#### Replicate #1

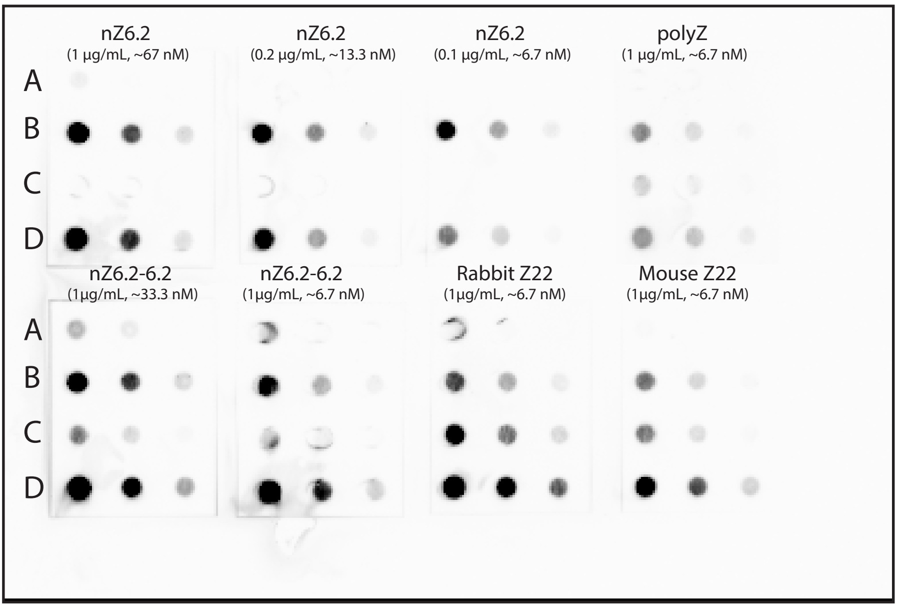


#### Replicate #2

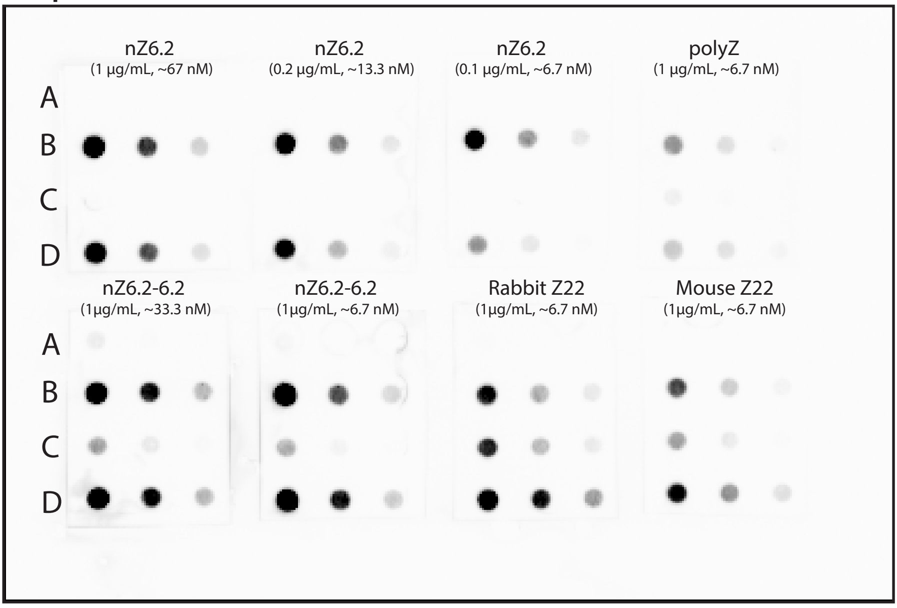


#### Replicate #3

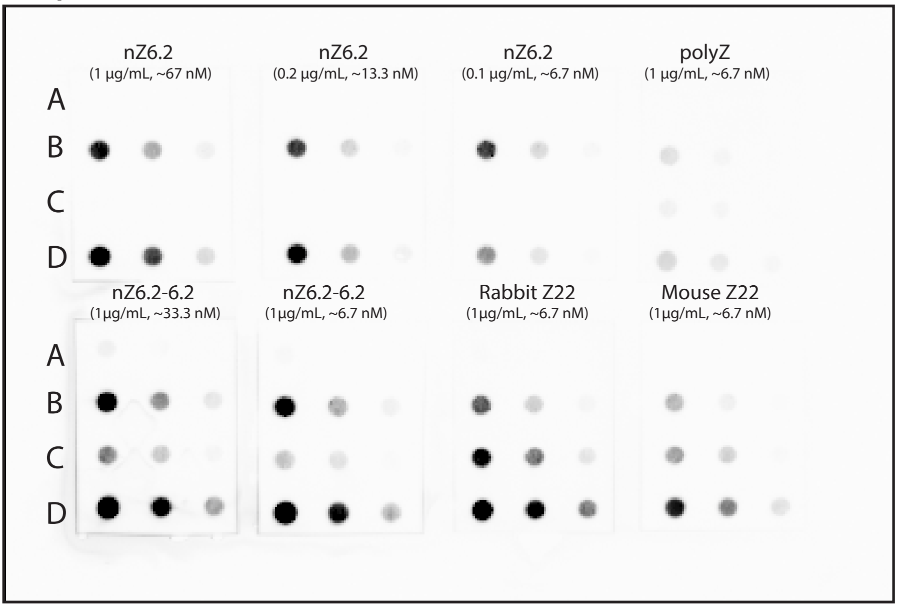
